# Human genetic analyses of organelles highlight the nucleus in age-related trait heritability

**DOI:** 10.1101/2021.01.22.427431

**Authors:** Rahul Gupta, Konrad J. Karczewski, Daniel Howrigan, Benjamin M. Neale, Vamsi K. Mootha

**Affiliations:** Howard Hughes Medical Institute and Department of Molecular Biology, Massachusetts General Hospital, Boston, MA 02114; Broad Institute of MIT and Harvard, Cambridge, MA 02142; Analytic and Translational Genetics Unit, Center for Genomic Medicine, Massachusetts General Hospital, Boston, MA 02114, USA

**Keywords:** mitochondria, nucleus, aging, oxidative phosphorylation, OXPHOS, transcription factor, zinc finger, KRAB domain, gene regulation, UK Biobank, constraint, dominance, haplosufficiency, haplosufficient, PPARGC1A, ESRRA, TFAM

## Abstract

Most age-related human diseases are accompanied by a decline in cellular organelle integrity, including impaired lysosomal proteostasis and defective mitochondrial oxidative phosphorylation. An open question, however, is the degree to which inherited variation in or near genes encoding each organelle contributes to age-related disease pathogenesis. Here, we evaluate if genetic loci encoding organelle proteomes confer greater-than-expected age-related disease risk. As mitochondrial dysfunction is a “hallmark” of aging, we begin by assessing nuclear and mitochondrial DNA loci near genes encoding the mitochondrial proteome and surprisingly observe a lack of enrichment across 24 age-related traits. Within nine other organelles, we find no enrichment with one exception: the nucleus, where enrichment emanates from nuclear transcription factors. In agreement, we find that genes encoding several organelles tend to be “haplosufficient,” while we observe strong purifying selection against heterozygous protein-truncating variants impacting the nucleus. Our work identifies common variation near transcription factors as having outsize influence on age-related trait risk, motivating future efforts to determine if and how this inherited variation then contributes to observed age-related organelle deterioration.

## Introduction

The global burden of age-related diseases such as type 2 diabetes (T2D), Parkinson’s disease (PD), and cardiovascular disease (CVD) has been steadily rising due in part to a progressively aging population. These diseases are often highly heritable: for example, narrow-sense heritabilities were recently estimated as 56% for T2D, 46% for general hypertension, and 41% for atherosclerosis^1^. Genome-wide association studies (GWAS) have led to the discovery of thousands of robust associations with common genetic variants^2^, implicating a complex genetic architecture as underlying much of the heritable risk. These loci hold the potential to reveal underlying mechanisms of disease and spotlight targetable pathways.

Aging has been associated with dysfunction in many cellular organelles^3^. Dysregulation of autophagic proteostasis, for which the lysosome is central, has been implicated in myriad age-related disorders including neurodegeneration, heart disease, and aging itself^4^, and mouse models deficient for autophagy in the central nervous system show neurodegeneration^5,6^. Endoplasmic reticular (ER) stress has been invoked as central to metabolic syndrome and insulin resistance in T2D^7^. Disruption in the nucleus through increased gene regulatory noise from epigenetic alterations^3^ and elevated nuclear envelope “leakiness”^8^ has been implicated in aging. Dysfunction in the mitochondria has even been invoked as a “hallmark” of aging^3^ and has been observed in many common age-associated diseases^9–15^. In particular, deficits in mitochondrial oxidative phosphorylation (OXPHOS) have been documented in aging and age-related diseases as evidenced by *in vivo* ^31^P-NMR measures^10,16^, enzymatic activity^11,12,17–21^ in biopsy material, accumulation of somatic mitochondrial DNA (mtDNA) mutations^13,14,22^, and a decline in mtDNA copy number (mtCN)^15^.

Given that a decline in organelle function is observed in age-related disease, a natural question is whether inherited variation in loci encoding organelles is enriched for age-related disease risk. Though it has long been known that recessive mutations leading to defects within many cellular organelles can lead to inherited syndromes (e.g., mutations in >300 nuclear DNA (nucDNA)-encoded mitochondrial genes lead to inborn mitochondrial disease^23^), it is unknown how this extends to common disease. In the present study, we use a human genetics approach to assess common variation in loci relevant to the function of ten cellular organelles. We begin with a deliberate focus on mitochondria given the depth of literature linking it to age-related disease, interrogating both nucDNA and mtDNA loci that contribute to the organelle’s proteome. This genetic approach is supported by the observation that heritability estimates of measures of mitochondrial function are substantial (33-65%^24,25^). We then extend our analyses to nine additional organelles.

To our surprise, we find no evidence of enrichment for genome-wide association signal in or near mitochondrial genes across any of our analyses. Further, of ten tested organelles, only the nucleus shows enrichment among many age-associated traits, with the signal emanating from the transcription factors (TFs). Further analysis shows that genes encoding the mitochondrial proteome tend to be tolerant to heterozygous predicted loss-of-function (pLoF) variation and thus are surprisingly “haplosufficient” – i.e., show little fitness cost with heterozygous pLoF. In contrast, nuclear TFs are especially sensitive to gene dosage and are often “haploinsufficient”, showing substantial purifying selection against heterozygous pLoF. Thus, our work highlights inherited variation influencing gene-regulatory pathways, rather than organelle physiology, in the inherited risk of common age-associated diseases.

## Results

### Age-related diseases and traits show diverse genetic architectures

To systematically define age-related diseases, we turned to recently published epidemiological data from the United Kingdom (U.K.)^26^ in order to match the U.K. Biobank (UKB)^27^ cohort. We prioritized traits whose prevalence increased as a function of age (**Methods**) and were represented in UKB (https://github.com/Nealelab/UK_Biobank_GWAS) and/or had available published GWAS meta-analyses^28–37^ (**Figure 1A, Supplementary note)**. We used SNP-heritability estimates from stratified linkage disequilibrium score regression (S-LDSC, https://github.com/bulik/ldsc)^38^ to ensure that our selected traits were sufficiently heritable (**Table S1, Methods, Supplementary note**), observing heritabilities across UKB and meta-analysis traits as high as 0.28 (bone mineral density), all with heritability Z-score > 4. We then computed pairwise genetic and phenotypic correlations between the age-associated traits to compare their respective genetic architectures and phenotypic relationships (**Figure 1B, Table S2, Methods**). In general, genetic correlations were greater in magnitude than respective phenotypic correlations, potentially as GWAS are less sensitive to purely non-genetic factors that influence phenotype (e.g., measurement error). As expected we find a highly correlated module of primarily cardiometabolic traits with high density lipoprotein (HDL) showing anti-correlation^39^. Interestingly, several other traits (gastroesophageal reflux disease (GERD), osteoarthritis) showed moderate genetic correlation to the cardiometabolic trait cluster while atrial fibrillation, for which T2D and CVD are risk factors^40^, showed phenotypic, but not genetic, correlation. Our final set of prioritized, age-associated traits included 24 genetically diverse, heritable phenotypes (**Table S1**). Of these, 11 traits were sufficiently heritable only in UKB, 3 were sufficiently heritable only among non-UKB meta-analyses, and 10 were well-powered in both UKB and an independent cohort.

**Figure 1.**
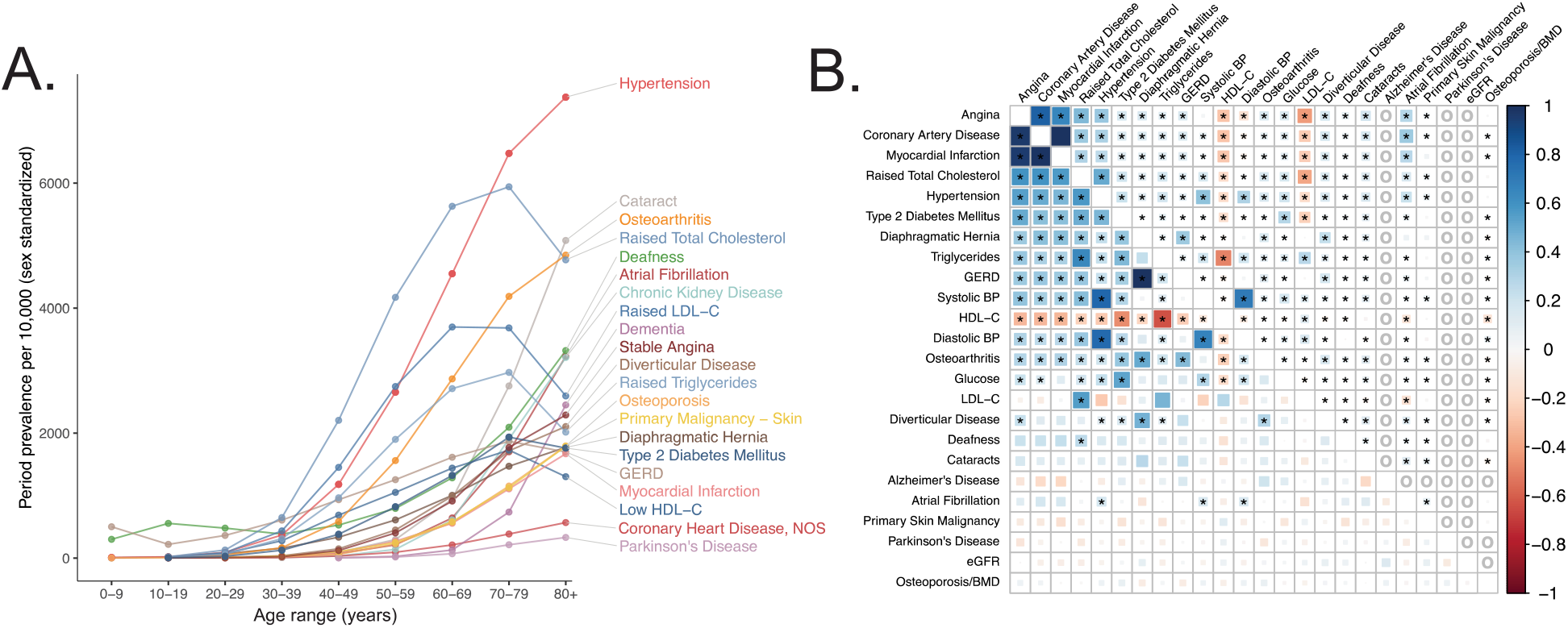
Selection of genetically diverse age-related diseases and traits using epidemiological data. **A**. Period prevalence of age-associated diseases systematically selected for this study **(Methods**). Epidemiological data obtained from Kuan et al. 2019. **B**: Genetic (lower half) and phenotypic (upper half) correlation between the selected age-related traits. All correlations were assessed between UK Biobank phenotypes with the exception of eGFR, Alzheimer’s Disease, and Parkinson’s Disease, for which the respective meta-analyses were used (**Methods**). Grey “o” in phenotypic correlations indicate phenotypes not tested within UKB for which individual-level data was not available. Point estimates and standard errors/p-values reported in **Table S2**. * represents correlations that are significantly different from 0 at a Bonferroni-corrected threshold for p = 0.05 across all tested traits.

### Mitochondrial genes are not enriched among age-related trait GWAS

To test if age-related trait heritability was enriched among mitochondria-relevant loci, we began by simply asking if ∼1100 nucDNA genes encoding the mitochondrial proteome from the MitoCarta2.0 inventory^41^ were found near lead SNPs for our selected traits represented in the NHGRI-EBI GWAS Catalog (https://www.ebi.ac.uk/gwas/)^42^ more frequently than expectation (**Methods, Supplementary note**). To our surprise, no traits showed a statistically significant enrichment of mitochondrial genes (**Figure 2-S1A**); in fact, six traits showed a statistically significant depletion. Even more strikingly, MitoCarta genes tended to be nominally enriched in fewer traits than the average randomly selected sample of protein-coding genes (**Figure 2-S1B**, empirical *p* = 0.014). This lack of enrichment was observed more broadly across virtually all traits represented in the GWAS Catalog (**Figure 2-S1C**). We also examined specific transcriptional regulators of mitochondrial biogenesis (*TFAM, GABPA, GABPB1, ESRRA, YY1, NRF1, PPARGC1A, PPARGC1B*) and found very little evidence supporting a role for these genes in modifying risk for the age-related GWAS Catalog phenotypes (**Supplementary note**).

**Figure 2.**
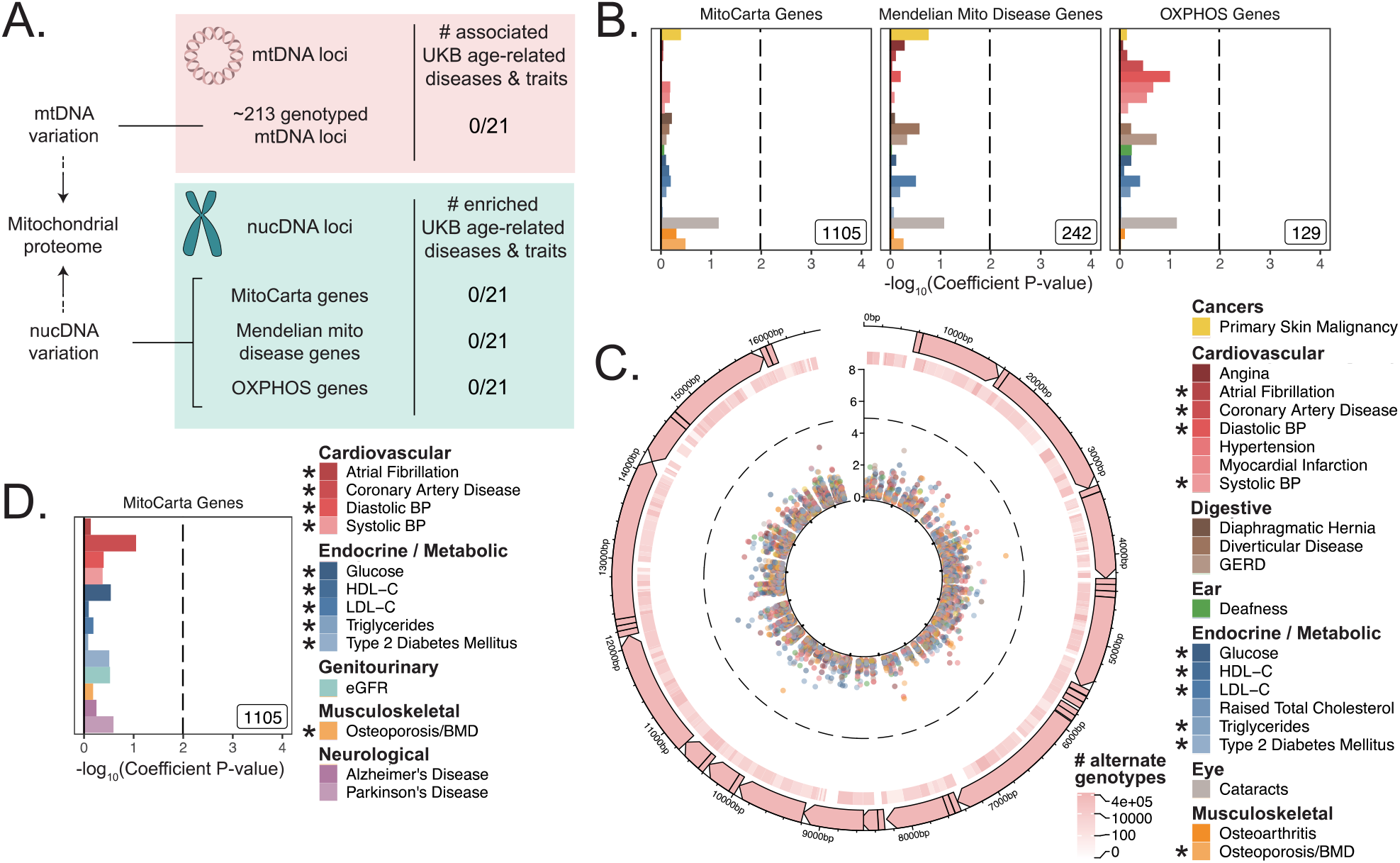
Assessment of the association of nucDNA and mtDNA loci contributing to the mitochondrial proteome with age-related traits. **A**. Scheme outlining the aspects of mitochondrial function assessed in this study. nucDNA loci contributing to the mitochondrial proteome are shown in teal, while mtDNA loci are shown in pink. **B**. S-LDSC enrichment p-values on top of the baseline model in UKB. Inset labels represent gene-set size; dotted line represents BH FDR 0.1 threshold. **C**. Visualization of mtDNA variants and associations with age-related diseases. The outer-most track represents the genetic architecture of the circular mtDNA. The heatmap track represents the number of individuals with alternate genotype on log scale. The inner track represents mitochondrial genome-wide association p-values, with radial angle corresponding to position on the mtDNA and magnitude representing –log_10_ P value. Dotted line represents Bonferroni cutoff for all tested trait-variant pairs. **D**. Replication of S-LDSC enrichment results in meta-analyses. Dotted line represents BH FDR 0.1 threshold. * represents traits for which sufficiently well powered cohorts from both UKB and meta-analyses were available. The trait color legend to the right of panel **C** applies to panels **B** and **C**, representing UKB traits.

To investigate further, we turned to U.K. Biobank (UKB). We compiled and tested loci encoding the mitochondrial proteome (**Figure 2A**) with which we interrogated the association between common mitochondrial variation and common disease. First, we considered all common variants in or near nucDNA MitoCarta genes, as well as two subsets of MitoCarta: mitochondrial Mendelian disease genes^23^ and nucDNA-encoded OXPHOS genes. Second, we obtained and tested mtDNA genotypes at up to 213 loci after quality control (**Methods**) from 360,662 individuals for associations with age-related traits.

First, we used S-LDSC^38,43^ and MAGMA (https://ctg.cncr.nl/software/magma)^44^, two robust methods that can be used to assess gene-based heritability enrichment accounting for LD and several confounders, to test if there was any evidence of heritability enrichment among MitoCarta genes (**Methods**). We found no evidence of enrichment near nucDNA MitoCarta genes for any trait tested in UKB using S-LDSC (**Figure 2B, 2-S7A**), consistent with our results from the GWAS Catalog. We replicated this lack of enrichment using MAGMA at two different window sizes (**Figure 2-S7C, 2-S7E;** all *q* > 0.1).

Given the lack of enrichment among the MitoCarta genes, we wanted to (1) verify that our selected methods could detect previously reported enrichments and (2) confirm that common variation in or near MitoCarta genes can lead to expression-level perturbations. We first successfully replicated previously reported enrichment among tissue-specific genes for key traits using both S-LDSC (**Figure 2-S2, 2-S3**) and MAGMA (**Figure 2-S4, 2-S5, Supplementary note, Methods**). We next confirmed that we had sufficient power using both S-LDSC and MAGMA to detect physiologically relevant enrichment effect sizes among MitoCarta genes (**Figure 2-S6, Methods, Supplementary note**). We finally examined the landscape of cis-expression QTLs (eQTLs) for these genes and found that almost all MitoCarta genes have cis-eQTLs in at least one tissue and often have cis-eQTLs in more tissues than most protein-coding genes (**Figure 2-S8, Methods, Supplementary note**). Hence, our selected methods could detect physiologically relevant heritability enrichments among our selected traits at gene-set sizes comparable to that of MitoCarta, and common variants in or near MitoCarta genes exerted *cis*-control on gene expression.

Next, we considered mtDNA loci genotyped in UKB, obtaining calls for up to 213 common variants passing quality control across 360,662 individuals (**Methods, Supplementary note**). We found no significant associations on the mtDNA for any of the 21 age-related traits available in UKB using linear or logistic regression (**Methods, Figure 2C, 2-S9, Table S4**).

As a control and to validate our approach, we also performed mtDNA-GWAS for specific traits with previously reported associations. A recent analysis of ∼147,437 individuals in BioBank Japan revealed four distinct traits with significant mtDNA associations^45^. Of these, creatinine and aspartate aminotransferase (AST) had sufficiently large sample sizes in UKB. We observed a large number of associations throughout the mtDNA for both traits (*p <* 1.15 * 10^−5^, **Figure 2-S9E**). Thus, our mtDNA association method was able to replicate robust mtDNA associations among well-powered traits.

We sought to replicate our negative results in an independent cohort. We turned to published GWAS meta-analyses^28–37^ (**Table S1**) and successfully replicated the lack of enrichment for MitoCarta genes across all 10 traits with an available independent cohort GWAS using S-LDSC (**Figure 2D, 2-S7B**) and MAGMA at two different window sizes (**Figure 2-S7D, Supplementary note;** all *q* > 0.1). Importantly, while we were unable to pursue analyses for PD and Alzheimer’s disease in UKB due to limited case counts, we tested MitoCarta genes among well-powered meta-analyses for these disorders (**Supplementary note**) and observed no enrichment (**Figure 2D;** all *q* > 0.1).

In summary, we tested (1) nucDNA loci near genes that encode the mitochondrial proteome in the GWAS Catalog, UKB, and GWAS meta-analyses, (2) transcriptional regulators of mitochondrial biogenesis in the GWAS Catalog, and (3) mtDNA variants in UKB. We found no convincing evidence of heritability enrichment for common age-associated diseases near these mitochondrial loci.

### Of all tested organelles, only the nucleus shows enrichment for age-related trait heritability

We next asked whether heritability for age-related diseases and traits clusters among loci associated with any cellular organelle. We used the COMPARTMENTS database (https://compartments.jensenlab.org) to define gene-sets corresponding to the proteomes of nine additional organelles^46^ besides mitochondria (**Methods**). We used S-LDSC to produce heritability estimates for these categories in the UKB age-related disease traits, finding evidence of heritability enrichment in many traits for genes comprising the nuclear proteome (**Figure 3A, Methods**). No other tested organelles showed evidence of heritability enrichment. Variation in or near genes comprising the nuclear proteome explained over 50% of disease heritability on average despite representing only ∼35% of tested SNPs (**Figure 3-S1, Supplementary note**). We successfully replicated this pattern of heritability enrichment among organelles using MAGMA in UKB at two window sizes (**Figure 3-S3A, 3-S3B**), again finding enrichment only among genes related to the nucleus.

**Figure 3.**
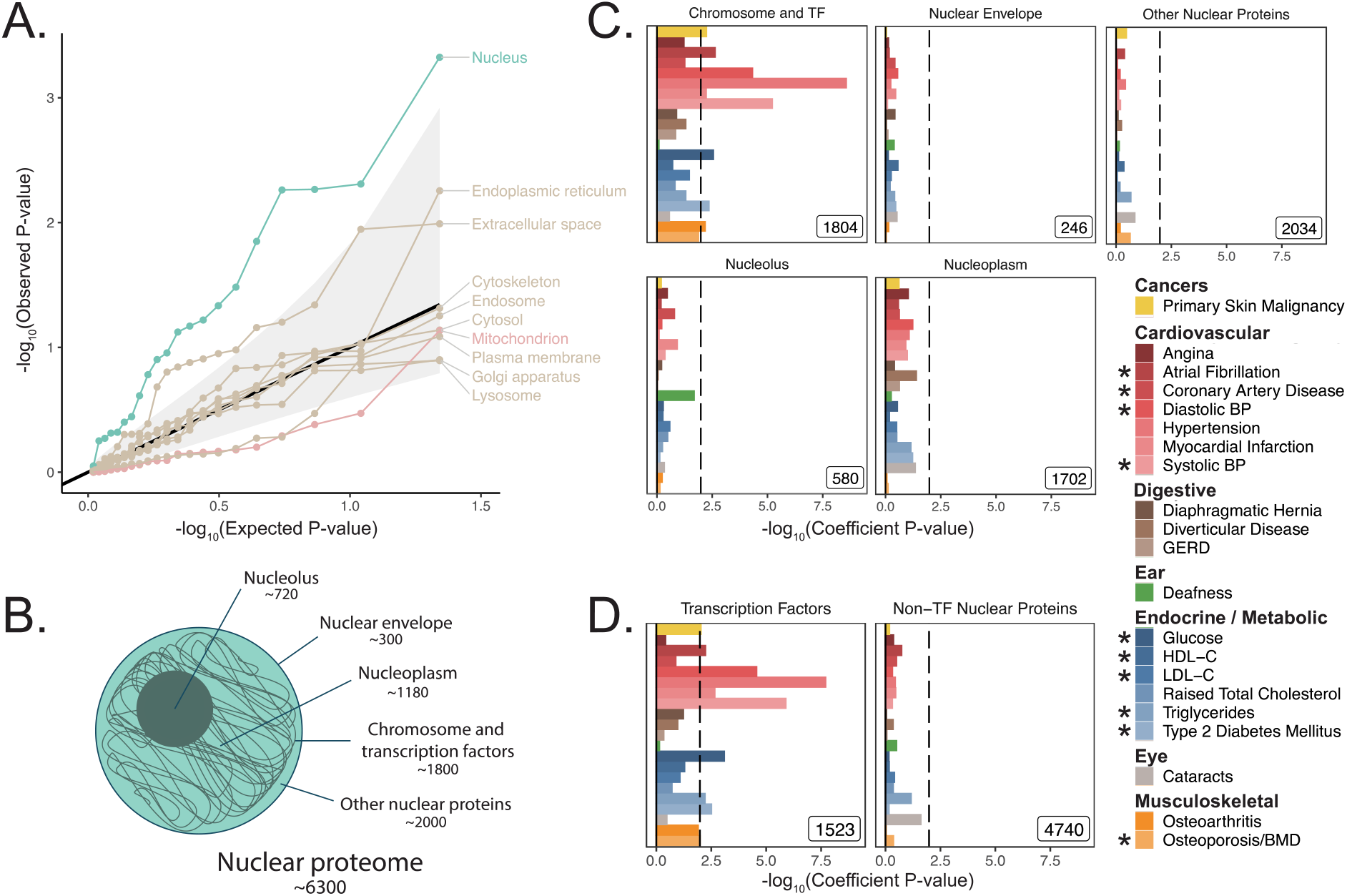
Heritability enrichment of organellar proteomes across age-related disease in UK Biobank. **A**. Quantile-quantile plot of heritability enrichment p-values atop the baseline model for gene-sets representing organellar proteomes, with black line representing expected null p-values following the uniform distribution and shaded ribbon representing 95% CI. **B**. Scheme of spatially distinct disjoint subsets of the nuclear proteome as a strategy to characterize observed enrichment of the nuclear proteome. Numbers represent gene-set size. **C**. S-LDSC enrichment p-values for spatial subsets of the nuclear proteome computed atop the baseline model. **D**. S-LDSC enrichment p-values for TFs and all other nucleus-localizing proteins. Inset numbers represent gene-set sizes, black lines represent cutoff at BH FDR < 10%. * represents traits for which sufficiently well powered cohorts from both UKB and meta-analyses were available.

### Much of the nuclear enrichment signal emanates from transcription factors

With over 6,000 genes comprising the nuclear proteome, we considered largely disjoint subsets of the organelle’s proteome to trace the source of the enrichment signal^47–49^ (**Figure 3B, Methods, Supplementary note**). We found significant heritability enrichment within the set of 1,804 genes whose protein products are annotated to localize to the chromosome itself (*q < 0*.*1* for 9 traits, **Figure 3C, 3-S2A**). Further partitioning revealed that much of this signal is attributable to the subset classified as TFs^49^ (1,523 genes, *q < 0*.*1* for 10 traits, **Figure 3D, 3-S2B**). We replicated these results using MAGMA in UKB at two window sizes (**Figure 3-S3**), and also replicated enrichments among TFs in several (but not all) corresponding meta-analyses (**Figure 3-S4**) despite reduced power (**Figure 2-S6H**). We generated functional subdivisions of the TFs (**Methods, Supplementary note**), finding that the non-zinc finger TFs showed enrichment for a highly similar set of traits to those enriched for the whole set of TFs (**Figure 3-S5D, 3-S6B, 3-S7B, 3-S8B**). Interestingly, the KRAB domain-containing zinc fingers (KRAB ZFs)^50^, which are recently evolved (**Figure 3-S5H**), were largely devoid of enrichment even compared to non-KRAB ZFs (**Figure 3-S5E, 3-S6C, 3-S7C, 3-S8C**). Thus, we find that variation within or near non-KRAB domain-containing TF genes has an outsize influence on age-associated disease heritability.

We next turned to recently published GWAS assessing parental lifespan^51^ and “healthspan” via first morbidity hazard^52^. Both traits showed highly significant heritability via S-LDSC (*h*^2^(*s. e*.) = 0.0265 (0.0019) and 0.0348 (0.003) respectively, **Methods**). Enrichment analysis of organelles among these traits revealed a significant enrichment for the nucleus for parental lifespan (*p* = 0.0003) using MAGMA (**Figure 4, Table S7**). While we observed only a nominally “suggestive” enrichment for the nucleus for healthspan (*p* = 0.058), S-LDSC showed significant nuclear heritability enrichment (*p* = 0.0016, **Figure 4-S1**). Analysis of spatial subsets of the nuclear proteome showed significant enrichment for TFs and proteins localizing to the chromosome in both aging phenotypes using MAGMA (**Figure 4**) and for healthspan using S-LDSC (**Figure 4-S1**).

**Figure 4.**
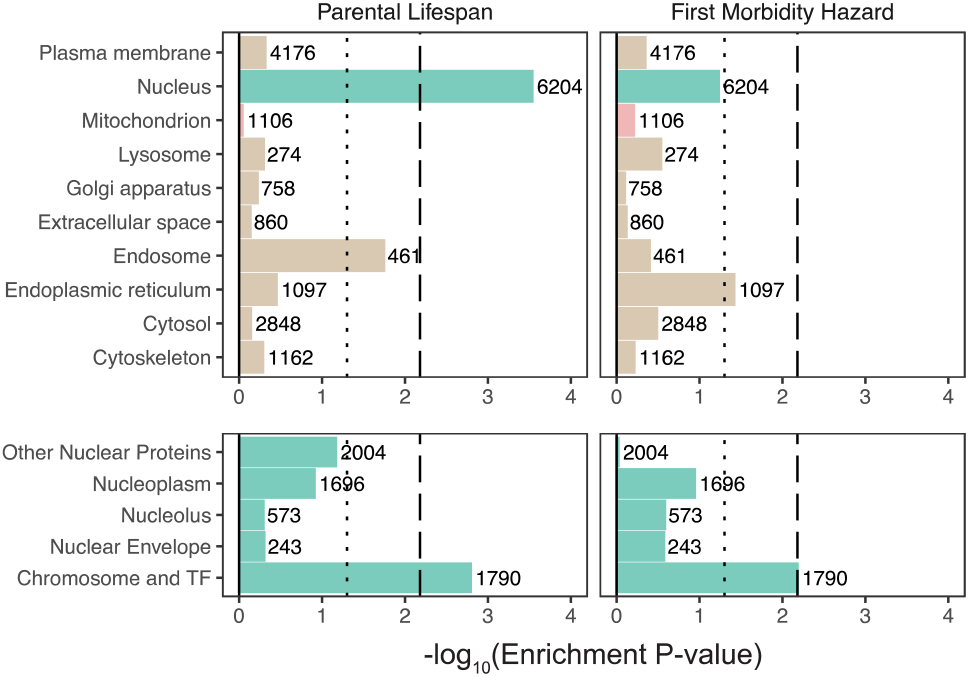
Enrichment of organellar proteomes within parental lifespan and healthspan as proxies for aging. Upper panels represent organelle proteomes; lower panels represent spatial subsets of the nuclear proteome. Numbers atop each bar represent gene-set sizes. Dashed lines represent cutoff at BH FDR < 10%, dotted lines represent nominal *p* = 0.05.

### Mitochondrial genes tend to be more “haplosufficient” than genes encoding other organelles

In light of observing heritability enrichment only among nuclear transcription factors, we wanted to determine if the fitness cost of pLoF variation in genes across cellular organelles mirrored our results. Mitochondria-localizing genes and TFs play a central role in numerous Mendelian diseases^23,53–55^, so we initially hypothesized that genes belonging to either category would be under significant purifying selection (i.e., constraint). We obtained constraint metrics from gnomAD (https://gnomad.broadinstitute.org)^56^ as the LoF observed/expected fraction (LOEUF). In agreement with our GWAS enrichment results, we observed that the mitochondrion on average is one of the least constrained organelles we tested, in stark contrast to the nucleus (**Figure 5A**). In fact, the nucleus was second only to the set of “haploinsufficient” genes (defined based on curated human clinical genetic data^56^, **Methods**) in the proportion of its genes in the most constrained decile, while the mitochondrion lay on the opposite end of the spectrum (**Figure 5B**). Interestingly, even the Mendelian mitochondrial disease genes had a high tolerance to pLoF variation on average in comparison to TFs (**Figure 5C**). Even across different categories of TFs, we observed that highly constrained TF subsets tend to show GWAS enrichment (**Figure 5-S1, 3-S5E**) relative to unconstrained subsets for our tested traits. Indeed, explicit inclusion of LOEUF as a covariate in the enrichment analysis model (**Methods**) reduced the significance of (but did not eliminate) the enrichment seen for the TFs (**Figure 5-S2B, 5-S3B, 5-S2E, 5-S2F**). Thus, while disruption in both mitochondrial genes and TFs can produce rare disease, the fitness cost of heterozygous variation in mitochondrial genes appears to be far lower than that among TFs. This dichotomy reflects the contrasting enrichment results between mitochondrial genes and TFs and supports the importance of gene regulation as it relates to evolutionary conservation.

**Figure 5.**
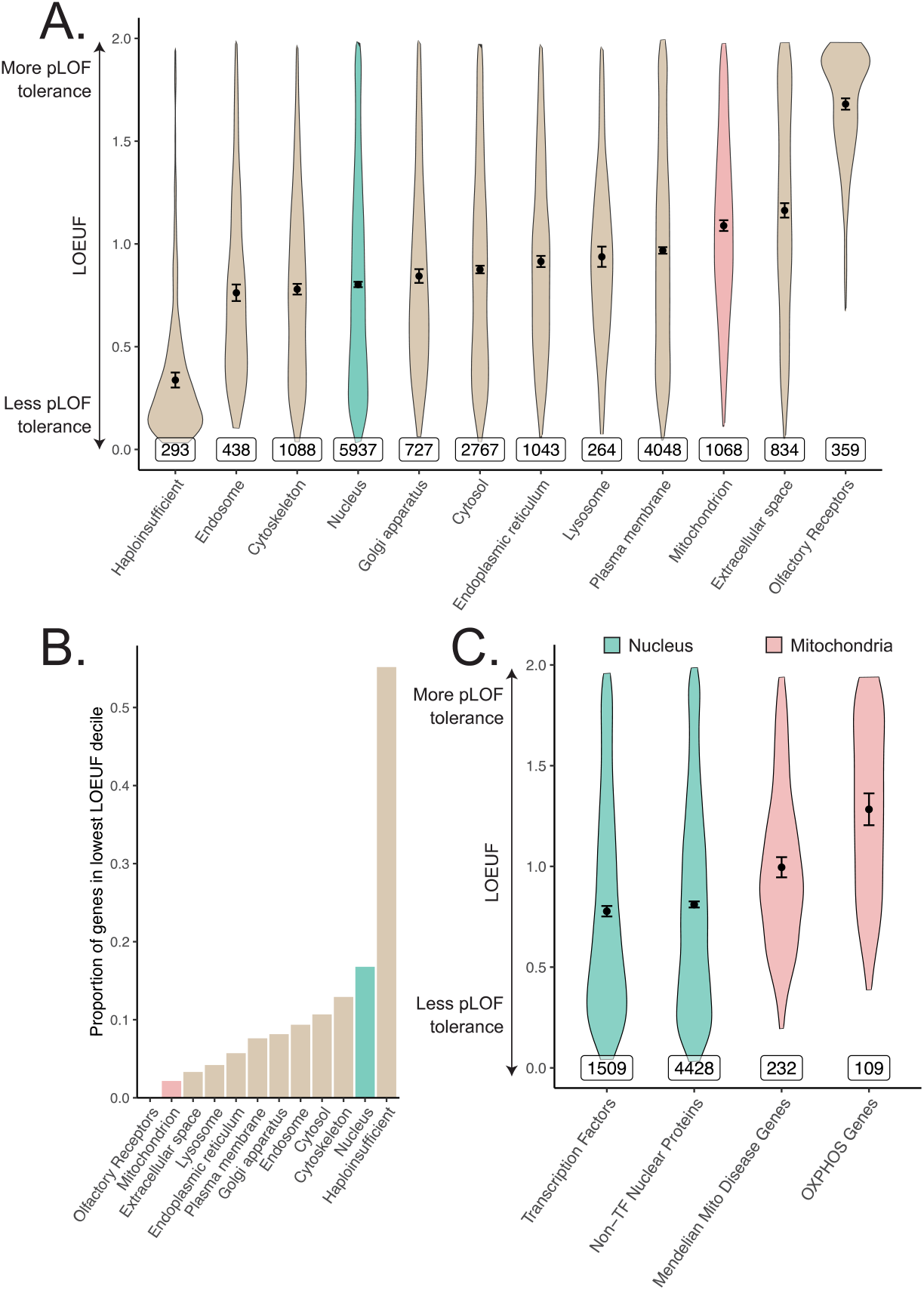
Differences in constraint distribution across organelles. **A**. Constraint as measured by LOEUF from gnomAD v2.1.1 for genes comprising organellar proteomes, book-ended by distributions for known haploinsufficient genes as well as olfactory receptors. Lower values indicate genes exacting a greater organismal fitness cost from a heterozygous LoF variant (greater constraint). **B**. Proportion of each gene-set found in the lowest LOEUF decile. Higher values indicate gene-sets containing more highly constrained genes. **C**. Constraint distributions for subsets of the nuclear-encoded mitochondrial proteome (red) and subsets of the nucleus (teal). Black points represent the mean with 95% CI. Inset numbers represent gene-set size.

## Discussion

Pathology in cellular organelles has been widely documented in age-related diseases^3,7,57–60^. Using a human genetics approach, here we report the unexpected discovery that except for the nucleus, cellular organelles tend not to be enriched in genetic associations for common, age-related diseases. We started with a focus on the mitochondria as a decline in mitochondrial abundance and activity has long been reported as one of the most consistent correlates of aging^14,16,18,22^ and age-associated diseases^10–13,15,17,19– 21^. We tested common variants contributing to the mitochondrial proteome on the nucDNA and mtDNA and found no convincing evidence of heritability enrichment in any tested trait, cohort, or method. We systematically expanded our analysis to survey 10 organelles and found that only the nucleus showed enrichment, with much of this signal originating from nuclear TFs. Constraint analysis showed a substantial fitness cost to heterozygous loss-of-function mutation in genes encoding the nuclear proteome, whereas genes encoding the mitochondrial proteome were “haplosufficient.”

Here, we focus on enrichment to place the complex genetic architectures of age-related traits in a broader biological context and prioritize pathways for follow-up. For these highly polygenic traits, any large fraction of the genome may explain a statistically significant amount of disease heritability^61,62^, and indeed associations between individual organelle-relevant loci and certain common diseases have been identified previously^63,64^. For example, variants in the endoplasmic reticular genes *WFS1* and *ATF6B* and the mitochondrial gene *ATP5G1* have been associated with common T2D^65^. These genes are present in the respective organelle gene-sets, however unlike TFs, neither the endoplasmic reticulum nor the mitochondrion showed enrichment for T2D. Importantly, both MAGMA and S-LDSC are capable of detecting an enrichment even in a highly polygenic background. Both methods have been used in the past to identify biologically plausible disease-relevant tissues^38,43^ and pathway enrichments^66,67^ in traits across the spectrum of polygenicity, and we identify enrichments among disease-relevant tissues using both methods in several highly polygenic traits.

While previous work has shown that common disease GWAS can be enriched for expression in specific disease-relevant organs^43,68^, our data suggest that this framework does not generally extend from organs to organelles. This finding contrasts with our classical nosology of inborn errors of metabolism that tend to be mapped to “causal” organelles, e.g., lysosomal storage diseases, disorders of peroxisomal biogenesis, and mitochondrial OXPHOS disorders. The observed enrichment for TFs within the nucleus indicates that common variation influencing genome regulation impacts common disease risk more than variation influencing individual organelles.

Our analysis of common inherited mitochondrial variation represents, to our knowledge, the most comprehensive joint assessment of mitochondria-relevant nucDNA and mtDNA variation in age-related diseases. We replicated mtDNA associations with creatinine and AST observed previously in BioBank Japan^45^, further supporting our approach. While individual mtDNA variants have been previously associated with certain traits^69–71^, these associations appear to be conflicting in the literature, perhaps because of limited power and/or uncontrolled confounding biases such as population stratification^72,73^. Our negative results are surprising, but they are compatible with a prior enrichment analysis focused on T2D^74^ as well as a small number of isolated reports interrogating either mitochondria-relevant nucDNA^74^ or mtDNA^45,75–77^ loci in select diseases.

To our knowledge, we are the first to systematically document heterogeneity in average pLoF across cellular organelles. That MitoCarta genes are “haplosufficient” and pLoF tolerant (**Figure 5A**) is consistent with the observation that most of the ∼300 inborn mitochondrial disease genes produce disease with recessive inheritance^23^ and healthy parents. The few mitochondrial disorders that show autosomal dominant inheritance are nearly always due to dominant negativity rather than haploinsufficiency. The intolerance of TFs to pLoF variation (**Figure 5C**) provide a stark contrast to the results from the mitochondria that is borne out in their associated Mendelian disease syndromes: TFs are known to be haploinsufficient^78^ and even regulatory variants modulating their expression can produce severe Mendelian disease^79^. We observe enrichment among TFs for 10 different diseases as well as parental lifespan and healthspan, consistent with observed elevated purifying selection against pLoF variants in these genes. Our enrichment results combined with pLoF intolerance suggest that variation among TFs may produce disease-associated variants with larger effect sizes than expectation, underscoring their importance as genetic “levers” for common disease heritability.

Why are mitochondria so robust to variation in gene dosage (**Figure 5**) and hence “haplosufficient?” We propose two possibilities. First, mitochondrial pathways tend to be highly interconnected, and it was already proposed by Wright^80^ and later by Kacser and Burns^81^ that haplosufficiency arises as a consequence of physiology, i.e., system output is inherently buffered against the partial loss of a single gene due to the network organization of metabolic reactions. Kacser and Burns in fact explicitly mention that noncatalytic gene products fall outside their framework, and we believe that our finding that nucleus-localizing and cytoskeletal genes are the two most pLoF-intolerant compartments is consistent with their assessment. Second, mitochondria were formerly autonomous microbes and hence may have retained vestigial layers of “intra-organelle buffering” against genetic variation. Numerous feedback control mechanisms, including respiratory control^82^, help to ensure organelle robustness across physiological extremes^83,84^. In fact, a recent CRISPR screen showed that of the genes for which knock-out modified survival under a mitochondrial poison, there is a striking over-representation of genes that themselves encode mitochondrial proteins^85^.

Throughout this study, we have tested for enrichment among inherited common variant associations near genes via an additive genetic model. We acknowledge the limitations of focusing on a specific genetic model and variant frequency regime, though note that common variation is the largest documented source of narrow-sense heritability, which typically accounts for a majority of disease heritability^86,87^. First, we consider only common variants. While rare variants may prove to be instructive, it is notable that a previous rare variant analysis in T2D^88^ failed to show enrichment among OXPHOS genes. Second, we consider only additive genetic models. A recessive model may be particularly fruitful for mitochondrial genes given their tolerance to pLoF variation, however these models are frequently power-limited and may not explain much more phenotypic variance than additive models^89,90^. Third, we have not considered epistasis. The effects of mtDNA-nucDNA interactions^91^ in common diseases have yet to be assessed. While there is debate about whether biologically-relevant epistasis can be simply captured by main effects^87,89,92,93^ at individual loci, it is possible that modeling mtDNA-nucDNA interactions will reveal new contributions. Fourth, to systematically assess all organelles, we restrict our analyses to variants near genes comprising each organelle’s proteome. It remains possible that future work will systematically identify novel organelle-relevant loci elsewhere in the genome which contribute disproportionately to age-related trait heritability. Fifth, while we are well-powered to detect physiologically relevant enrichments among most tested organelles (including the mitochondrion), our power may be more limited for particularly small compartments (e.g., lysosome). Finally, it is crucial not to confuse our mtDNA-GWAS results with previously reported associations between somatic mtDNA mutations and age-associated disease^13,14,22^ – the present work is focused on germline variation.

We have not formally addressed the causality of mitochondrial dysfunction in common age-related disease and the observed lack of heritability enrichment does not preclude the possibility of a therapeutic benefit in targeting the mitochondrion for age-related disease. For example, mitochondrial dysfunction is documented in brain or heart infarcts following blood vessel occlusion in laboratory-based models^94,95^. Clearly mitochondrial genetic variants do not influence infarct risk in this laboratory model, but pharmacological blockade of the mitochondrial permeability transition pore can mitigate reperfusion injury and infarct size^96^. Future studies will be required to determine if and how the mitochondrial dysfunction associated with common age-associated diseases can be targeted for therapeutic benefit. Efforts to develop reliable measures of mitochondrial function and dysfunction have the potential to unbiasedly discover genetic instruments that influence the mitochondrion, and causal inference techniques such as Mendelian Randomization may shed light on this important causal question.

Our finding that the nucleus is the only organelle that shows enrichment for common age-associated trait heritability builds on prior work implicating nuclear processes in aging. Most human progeroid syndromes result from monogenic defects in nuclear components^97^ (e.g., *LMNA* in Hutchinson-Gilford progeria syndrome, *TERC* in dyskeratosis congenita), and telomere length has long been observed as a marker of aging^98^. Heritability enrichment of age-related traits among gene regulators is consistent with the epigenetic dysregulation^99^ and elevated transcriptional noise^3,100^ observed in aging (e.g., *SIRT6* modulation influences mouse longevity^101^ and metabolic syndrome^58^). An important role for gene regulation in common age-related disease is in agreement with both the observation that a very large fraction of common disease-associated loci corresponds to the non-coding genome and the enrichment of disease heritability in histone marks and TF binding sites^38,102^. Given that a deterioration in several other cellular organelles has been so frequently documented in age-related traits, a future challenge lies in elucidating how inherited variation in or near TFs ultimately leads to the observed organelle dysfunction in age-related disease.

## Acknowledgements

We thank D. Altshuler, S.E. Calvo, T. Finkel, H. Finucane, E.S. Lander, M.E. MacDonald, D. Palmer, E.B. Robinson, A.V. Segrè, M.E. Talkowski, R.K. Walters, C.C. Winter, and members of the Mootha and Neale labs for critical feedback and discussions. This research has been conducted using the UK Biobank Resource under Application Number 31063. This project was supported in part by grants (NIH R35GM122455 to V.K.M. and NIH T32 AG000222 to R.G.) from the National Institutes of Health. VKM is an Investigator of the Howard Hughes Medical Institute.

## Data Availability

Heritability point estimates and standard errors for age-related traits are listed in Table S1. Genetic and phenotypic correlation point estimates and standard errors/p-values plotted in Figure 1B are available in Table S2. Summary statistics from mtDNA-GWAS are available in Table S4. All gene-based enrichment analysis p-values and point estimates are available in Tables S6 and S7. Period prevalence data for diseases in the UK can be obtained from Kuan et al. 2019. Gene-sets can be found using COMPARTMENTS (https://compartments.jensenlab.org), MitoCarta 2.0 (https://www.broadinstitute.org/files/shared/metabolism/mitocarta/human.mitocarta2.0.html), Lambert et al. 2018 (DOI: 10.1016/j.cell.2018.01.029), Frazier et al. 2019 (DOI: 10.1074/jbc.R117.809194), Finucane et al. 2018 (https://alkesgroup.broadinstitute.org/LDSCORE/), Kapopoulou et al. 2015 (DOI: 10.1111/evo.12819), and the Macarthur laboratory (https://github.com/macarthur-lab/gene_lists). Gene age estimates were obtained from Litman, Stein 2019 (DOI: 10.1053/j.seminoncol.2018.11.002). GWAS catalog annotations can be obtained from: https://www.ebi.ac.uk/gwas. Heritability estimates across UKB can be obtained at: https://nealelab.github.io/UKBB_ldsc/. UKB summary statistics can be obtained from Neale lab GWAS round 2: https://github.com/Nealelab/UK_Biobank_GWAS. Annotations for the Baseline v1.1 and BaselineLD v2.2 models as well as other relevant reference data, including the 1000G EUR reference panel, can be obtained from https://alkesgroup.broadinstitute.org/LDSCORE/. eQTL and expression data in human tissues can be obtained from GTEx (https://www.gtexportal.org). Constraint estimates can be found via gnomAD: https://gnomad.broadinstitute.org. See citations for publicly available GWAS meta-analysis summary statistics^28–37,51,52^.

## Code Availability

Our analysis leverages publicly available tools including LDSC for heritability enrichment and genetic correlation (https://github.com/bulik/ldsc), MAGMA v1.07b for gene-set enrichment analysis (https://ctg.cncr.nl/software/magma), and Hail v0.2.51 for distributed computing and mtDNA GWAS (https://hail.is).

## Author Contributions

R.G., B.M.N., and V.K.M. conceived of the project; R.G., K.J.K., D.H. designed analyses; R.G. performed analyses; B.M.N., V.K.M. supervised project; R.G. and V.K.M. wrote the manuscript with input from other authors.

## Materials and Methods

### Trait selection

Sex-standardized period prevalence of over 300 diseases was obtained from an extensive survey of the National Health Service in the UK as reported previously^26^. To select high prevalence late-onset diseases, we ranked diseases with a median onset over 50 years of age by the sum of the period prevalence of all age categories above 50. We selected the top 30 diseases using this metric and manually mapped these traits to similar or equivalent phenotypes with publicly available summary statistics from UKB and/or well-powered meta-analyses (e.g., Parkinson’s Disease and Alzheimer’s Disease for dementia) resulting in 24 traits with data available in UKB, meta-analyses, or both (**Table S1**).

### Criteria for inclusion of summary statistics

We manually mapped selected age-related diseases and traits to corresponding phenotypes in UKB. In parallel, we searched the literature to identify well-powered EUR-predominant GWAS (referred to as meta-analyses) that (1) used primarily non-targeted arrays, (2) had publicly available full summary statistics, and (3) did not enroll individuals from UKB to serve as independent replication (**Supplementary note**). We produced heritability estimates using stratified linkage-disequilibrium score regression (S-LDSC, https://github.com/bulik/ldsc)^38^ atop the BaselineLD v2.2 model using reference LD scores computed from 1000G EUR (https://alkesgroup.broadinstitute.org/LDSCORE/). We computed the heritability Z-score, a statistic that captures sample size, polygenicity, and heritability^38^, and included only traits with heritability Z-score > 4 (**Supplementary note**) for further analysis.

### Genetic correlations among age-related traits

Pairwise genetic correlations, *r*_*g*_, were computed using linkage-disequilibrium score correlation^39^ on all selected age-related traits with heritability Z-score > 4. We used UKB summary statistics (https://github.com/Nealelab/UK_Biobank_GWAS) for all sufficiently powered traits; summary statistics from meta-analyses were used for eGFR^35^, Alzheimer’s Disease^37^, and Parkinson’s Disease^36^ as these traits showed heritability Z-score > 4 within meta-analyses but not in UKB **(Table S1)**. P-values for genetic correlation represented deviation from the null hypothesis *r*_*g*_ = 0. Traits were ordered by their contribution to the first eigenvector of the absolute value of the correlation matrix, with point estimates and standard errors available in **Table S2**. Bonferroni correction was applied producing a p-value cutoff of 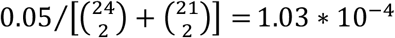, accounting for both genotypic and phenotypic correlation hypothesis tests.

### Phenotypic correlations in UKB

Pairwise phenotypic correlations, *r*_*p*_, were computed for all 21 traits with well-powered individual level data available in UKB (**Table S1**). Pearson correlation was computed between continuous traits via `cor.test` in R with a two-sided alternative. Tetrachoric correlation was used to compute correlations between binary traits and biserial correlation was used for correlations between binary and continuous traits, using the `polychor` and `polyserial` functions of the `polycor` package in R respectively using the two-step approximation. These approaches model a latent normally distributed variable underlying binary traits. P-values were computed using a normal approximation using standard error estimates from `polycor`. Point estimates and standard errors are available in **Table S2**.

### Assessment of mitochondria-localizing genes in the GWAS Catalog

We mapped variants in the GWAS Catalog (obtained on September 5^th^, 2019, https://www.ebi.ac.uk/gwas/) meeting genome-wide significance (p < 5e-8) to genes using provided annotations, producing a set of trait-associated genes for each trait. We manually selected phenotypes represented in the GWAS Catalog matching our set of age-associated traits with > 30 trait-associated genes. For each trait, we computed the proportion of trait-associated genes that were mitochondria-localizing (defined via MitoCarta2.0^41^) and tested for enrichment or depletion relative to overall genome background using two-sided Fisher’s exact tests correcting for multiple hypothesis tests with the Benjamini-Hochberg (BH) procedure at FDR q-value < 0.1.

We also computed the test statistic 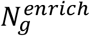, defined as the number of age-associated traits showing a nominal (not necessarily statistically significant) enrichment for a given gene-set *g*, for the MitoCarta genes. We then generated an empirical null distribution for 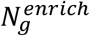. We drew 1,000 random samples of protein-coding genes, where each sample contained the same number of genes as the set of mitochondria-localizing genes and computed 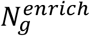 for each of these gene-sets **(Figure 2-S1B)**. The one-sided p-value, defined as 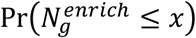 under the null, was subsequently obtained.

We expanded our enrichment/depletion analysis to all 332 traits in the GWAS Catalog with over 30 trait-associated genes; for enrichment or depletion testing, we used two-sided Fisher’s exact tests and corrected for multiple hypothesis testing with the BH procedure at FDR q-value < 0.1.

### Harmonization and filtering of summary statistics for LDSC and MAGMA

UKB summary statistics previously formatted for use with LDSC and filtered to HapMap3 (HM3) SNPs (https://github.com/Nealelab/UKBB_ldsc) were used for analysis with S-LDSC. For analysis with MAGMA v1.07b^44^, we included variants from the full Neale Lab UKB Round 2 GWAS summary statistics (https://github.com/Nealelab/UK_Biobank_GWAS) with INFO > 0.8 and MAF > 0.01, and excluded any variants flagged as low confidence (a heuristic defined by MAF < 0.001 or expected case MAC < 25).

Summary statistics obtained from publicly available GWAS meta-analyses^28–37^ were reported in varied formats. We manually verified the genome build upon which each meta-analysis reported results and ensured that all sets of summary statistics contained columns listing P-value, variant rsID, genome-build specific coordinates, and if available, variant-specific sample size (**Table S1**). If variant coordinates or rsID were not provided, the relevant columns were obtained from dbSNP database version 130 (for hg18) or 146 (for hg19). We used the summary statistic munging script provided with S-LDSC (https://github.com/bulik/ldsc) to generate summary statistics compatible with S-LDSC, restricting to HM3 SNPs as these tend to be best behaved for analysis with LDSC. For use of meta-analyses with MAGMA^44^, we restricted analysis to variants with INFO > 0.8 and MAF > 0.01 if such information was provided.

### Multiple testing correction for gene-set enrichment analysis

To account for the multiple hypothesis tests performed throughout this study for age-related traits, we obtained p-value thresholds via the BH procedure at FDR < 0.1 for all gene-sets assessed for a given method and cohort type (where the two cohort types were UKB and meta-analysis). The BH procedure at FDR < 0.1 was also applied to our analyses of parental lifespan and healthspan.

### Gene-set based enrichment analysis

We extensively use S-LDSC and MAGMA to perform gene-set enrichment analyses among GWAS summary statistics. To test enrichment with S-LDSC, SNPs were mapped to each gene with a 100kb symmetric window as recommended^43^ and LD scores were computed using the 1000G EUR reference panel (https://alkesgroup.broadinstitute.org/LDSCORE/) and subsequently restricted to the HM3 SNPs. We used S-LDSC to test for heritability enrichment controlling for 53 annotations including coding regions, enhancer regions, 5’ and 3’ UTRs, and others as previously described^38^ (baseline v1.1, referred to as baseline model hereafter). We also used MAGMA with both 5kb up, 1.5kb down and 100kb symmetric windows to test for enrichment. MAGMA gene-level analysis was performed with the 1000G EUR LD reference panel to account for LD structure, and gene-set analysis was performed including covariates for gene length, variant density, inverse minor allele count (MAC), as well as log-transformed versions of these covariates. Statistical tests for both S-LDSC and MAGMA were one-sided, considering enrichment only. For both methods, we included the relevant superset of genes as a control to ensure that our analysis was competitive (**Supplementary note**). We refer to this approach as the ‘usual approach’. All enrichment effect size estimates and p-values are available in **Tables S6** and **Table S7**.

### Enrichment analysis of genes comprising the mitochondrial proteome

We obtained the set of nuclear-encoded mitochondria-localizing genes using MitoCarta2.0^41^ and used the literature to obtain the subset of MitoCarta genes involved in inherited mitochondrial disease^23^ as well as those producing components of oxidative phosphorylation (OXPHOS) complexes. We used both S-LDSC and MAGMA to test for enrichment in the usual way (**Methods**) controlling for the set of protein-coding genes to ensure a competitive analysis (**Supplementary note**). We also tested mitochondria-localizing genes for enrichment in meta-analyses using S-LDSC and MAGMA with the same parameters as for UKB traits (**Supplementary note**).

### Tissue-expressed gene-set enrichment analysis

To obtain the set of genes most expressed in a given tissue versus others, we obtained t-statistics computed from GTEx v6 gene-level transcript-per-million (TPM) data corrected for age and sex as published previously^43^. For each tissue, we selected the top 2485 genes (10%) with the highest t-statistics for tissue-specific expression, producing tissue-expressed gene-sets. We selected nine tissues based on expectation of enrichment for our tested traits in UKB (e.g., liver for LDL levels, esophageal mucosa for GERD). We used both S-LDSC and MAGMA to test for enrichment in the usual way (**Methods**) controlling for the set of tissue-expressed genes to ensure a competitive analysis (**Supplementary note**). Tissue-expressed gene-set analyses were performed on meta-analyses with S-LDSC and MAGMA on the same tissues using the same parameters as used in UKB.

### Power analysis

To test for the effects of gene-set size on power, we selected ten positive control tissue-trait pairs based on (1) the presence of tissue enrichment in UKB with S-LDSC and MAGMA and (2) if the observed enrichment was biologically plausible. The pairs tested were liver-HDL, liver-LDL, liver-TG, liver-cholesterol, pancreas-glucose, pancreas-T2D, atrial appendage-atrial fibrillation, sigmoid colon-diverticular disease, coronary artery-myocardial infarction, and visceral adipose-HDL. We then, in brief, used an empirical sampling-based approach, generating random subsamples of a selected set of tissue-expressed gene-sets at four different gene-set sizes (1523, 1105, 800, and 350 genes), defining power as the proportion of trials showing a significant enrichment (**Supplementary note**). We used the same sub-sampled gene-sets for enrichment analysis using both S-LDSC and MAGMA in the usual way (**Methods**) controlling for the set of tissue-expressed genes to ensure a competitive analysis (**Supplementary note**). We used the same gene-sets among the subset of the positive control traits that showed enrichment in the corresponding meta-analysis to verify power for the meta-analyses (**Supplementary note**).

### Cross-tissue eQTL analysis

We obtained the set of eGenes from GTEx v8 across 49 tissues (https://www.gtexportal.org), filtering to only include cis-eQTLs with q-value < 0.05. To determine how the landscape of cis-eQTLs for MitoCarta genes compared to other protein-coding genes, we regressed the number of tissues with a detected cis-eQTL for a given gene x, 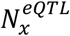, onto an indicator for membership in a given organellar proteome 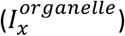, controlling for gene length, log gene length, breadth of expression (τ.), and the number of tissues with detected expression > 5 TPM (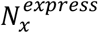, **Supplementary note**). To quantify breadth of expression, we obtained median-per-tissue GTEx v8 TPM expression values and computed τ^103^ after removing lowly-expressed genes with maximal cross-tissue TPM < 1, defined as:

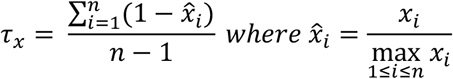

where *x*_*i*_ is the expression of gene *x* in tissue *i* with *n* tissues. τ ranges from 0 to 1, with lower τ indicating broadly expressed gene and higher τ indicating more tissue specific expression patterns. Because GTEx sampled multiple tissue subtypes (e.g., brain sub-regions) that show correlated expression profiles^104^ which bias *τ*_*x*_, 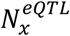, and 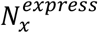 upward, for each broader tissue class (brain, heart, artery, esophagus, skin, cervix, colon, adipose) we selected a single representative tissue when computing these quantities (**Figure 3-S5B, Supplementary note**). We used LD scores computed from the 1000G EUR reference panel. The model, fit via OLS for each tested organelle, was:

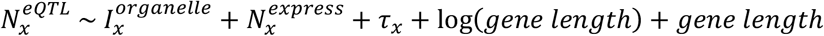

### mtDNA-wide association study

We obtained mtDNA genotype data on 265 variants as obtained on the UK Biobank Axiom array and the UK BiLEVE array from the full UKB release^27^. To perform variant QC, we used evoker-lite^105^ to generate fluorescence cluster plots per-variant and per-batch and manually inspected the results, removing 19 variants due to cluster plot abnormalities (**Table S3, Supplementary note**). We additionally removed any variants with heterozygous calls, within-array-type call rate < 0.95, and with less than 20 individuals with an alternate genotype. For case-control traits, we removed any phenotype-variant pair with an expected case count of alternate genotype individuals of less than 20, resulting in a maximum of 213 variants tested per trait (**Supplementary note**). To perform sample QC, we restricted samples to the same samples from which UKB summary statistics were generated (https://github.com/Nealelab/UK_Biobank_GWAS), namely unrelated individuals 7 standard deviations away from the first 6 European sample selection PCs with self-reported white-British, Irish, or White ethnicity and no evidence of sex chromosome aneuploidy. We additionally removed any samples with within-array-type mitochondrial variant call rate < 0.95, resulting in 360,662 unrelated samples of EUR ancestry. We generated the LD matrix for mitochondrial DNA variants using Hail v0.2.51 (https://hail.is) pairwise for all 213 variants tested across all post-QC samples.

We ran mtDNA-GWAS for all 21 UKB age-related phenotypes as well as creatinine and AST using Hail v0.2.51 via linear regression controlling for the first 20 PCs of the nuclear genotype matrix, sex, age, age^2^, sex*age, and sex*age^2^ as performed for the UKB GWAS (https://github.com/Nealelab/UK_Biobank_GWAS). We also used Hail to run Firth logistic regression with the same covariates for case/control traits. As we observed that some mitochondrial DNA variants were specific to array type, we also ran linear regression including array type as a covariate; we did not perform logistic regression with array type as a covariate due to convergence issues from complete separation of variants assessed only on a single array type. We defined mtDNA-wide significance using a Bonferroni correction by 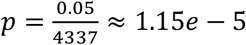.

### Enrichment analysis of components of organellar proteomes

COMPARTMENTS (https://compartments.jensenlab.org)^46^ is a resource integrating several lines of evidence for protein localization predictions including annotations, text-mining, sequence predictions, and experimental data from the Human Protein Atlas. We used this resource to obtain the degree of evidence (a number ranging from 0 to 5) linking each gene to localization to one of 12 organelles: nucleus, cytosol, cytoskeleton, peroxisome, lysosome, endoplasmic reticulum, Golgi apparatus, plasma membrane, endosome, extracellular space, mitochondrion, and proteasome. To avoid noisy localization assignments due to weak text mining and prediction evidence, we only considered localization assignments with a score > 2 as described previously^46^. We subsequently assigned compartment(s) to each gene by selecting the compartment(s) with the maximal score within each gene. We only included compartments containing over 240 genes due to limited power at smaller gene-set sizes and used MitoCarta2.0^41^ to obtain a higher confidence set of genes localizing to the mitochondrion, resulting in gene-sets representing the proteomes of 10 organelles. S-LDSC and MAGMA were used to test for enrichment across the UKB age-related traits for these gene-sets in the usual way, controlling for the set of protein-coding genes. S-LDSC was also used to obtain estimates of the percentage of heritability explained by each organelle gene-set.

### Enrichment analysis of spatial components of the nucleus

To produce interpretable sub-divisions of the nucleus, we used Gene Ontology (GO)^47,48^ to identify terms listed as children of the nucleus cellular component (GO:0005634). We used Ensembl version 99^106^ to obtain a first pass set of genes annotated to each sub-compartment of the nucleus (or its children). After manual review of sub-compartments with > 90 genes, we selected nucleoplasm (GO:0005654), nuclear chromosome (GO:0000228), nucleolus (GO:0005730), nuclear envelope (GO:0005635), splicosomal complex (GO:0005681), nuclear DNA-directed RNA polymerase complex (GO:0055029), and nuclear pore (GO:0005643). We excluded terms listed as ‘part’ due to poor interpretability and manually excluded similar terms (e.g., nuclear lumen vs nucleoplasm). To generate a high confidence set of genes localizing to each of these selected sub-compartments, we then turned to the COMPARTMENTS resource which assigns localization confidence scores for each protein to GO cellular component terms. We assigned members of the nuclear proteome to these selected nuclear sub-compartments using same the approach outlined for the organelle analysis (**Methods**). After filtering our selected sub-compartments to those containing > 240 genes, we obtained four categories: nucleoplasm, nuclear chromosome, nucleolus, and nuclear envelope. The nuclear chromosome annotation was largely overlapping with a manually curated high-quality list of TFs^49^ however was not exhaustive; as such, we merged these lists to generate the chromosome and TF category. To improve interpretability, we removed genes from nucleoplasm that were also assigned to another nuclear sub-compartment, constructed a list of other nucleus-localizing proteins not captured in these four sub-compartments, and included only genes annotated as localizing to the nucleus (**Methods**). S-LDSC and MAGMA were used to test for enrichment across the UKB age-related traits for these gene-sets in the usual way while controlling for the set of protein-coding genes (**Methods**).

### Enrichment analysis of functionally distinct TF subsets

We used a published curated high-quality list of TFs^49^ to partition the Chromosome and TF category into TFs and other chromosomal proteins. To determine which TFs are broadly expressed versus tissue specific, we computed τ per TF across all selected tissues after removing lowly-expressed genes with maximal cross-tissue TPM < 1 (**Methods, Supplementary note**). The threshold for tissue-specific genes was set at τ ≥ 0.*76* based on the location of the central nadir of the resultant bimodal distribution (**Figure 3-S5A**). To identify terciles of TFs by age, we obtained relative gene age assignments for each gene previously generated by obtaining the modal earliest ortholog level across several databases mapped to 19 ordered phylostrata^107^. DNA binding domain (DBD) annotations for the TFs were obtained from previous manual curation efforts^49^. S-LDSC and MAGMA were used to test for enrichment across the UKB age-related traits for these gene-sets in the usual way while controlling for the set of protein-coding genes (**Methods**). We also tested TFs for enrichment in meta-analyses using S-LDSC and MAGMA with the same parameters as for UKB traits (**Supplementary note**).

### Analysis of constraint across organelles and sub-organellar gene-sets

We obtained gene-level gnomAD v2.1.1 constraint tables (https://gnomad.broadinstitute.org), haploinsufficient genes, and olfactory receptors^56^ (https://github.com/macarthur-lab/gene_lists). Constraint values as loss-of-function observed/expected fraction (LOEUF) were mapped to genes within organelle, sub-mitochondrial, sub-nuclear, and TF binding domain gene-sets.

### Enrichment analysis across age-related disease holding constraint as a covariate

To test for enrichment with constraint as a covariate, we used MAGMA with UKB age-related traits. We mapped variants to genes and performed the gene-level analysis as done previously for the mitochondria-localizing gene and organelle analysis. We included LOEUF and log LOEUF as covariates for the gene-set analysis in addition to the default covariates (gene length, SNP density, inverse MAC, as well as the respective log-transformed versions) via the `–condition-residualize` flag.

## References

1. Wang K, Gaitsch H, Poon H, Cox NJ, Rzhetsky A. Classification of common human diseases derived from shared genetic and environmental determinants. Nat Genet. 2017;49(9):1319–1325. doi:10.1038/ng.3931

2. Claussnitzer M, Cho JH, Collins R, et al. A brief history of human disease genetics. Nature. 2020;577(7789):179–189. doi:10.1038/s41586-019-1879-7

3. López-Otín C, Blasco MA, Partridge L, Serrano M, Kroemer G. The hallmarks of aging. Cell. 2013;153(6):1194. doi:10.1016/j.cell.2013.05.039

4. Mizushima N, Levine B, Cuervo AM, Klionsky DJ. Autophagy fights disease through cellular self-digestion. Nature. 2008;451(7182):1069–1075. doi:10.1038/nature06639

5. Hara T, Nakamura K, Matsui M, et al. Suppression of basal autophagy in neural cells causes neurodegenerative disease in mice. Nature. 2006;441(7095):885–889. doi:10.1038/nature04724

6. Komatsu M, Waguri S, Chiba T, et al. Loss of autophagy in the central nervous system causes neurodegeneration in mice. Nature. 2006;441(7095):880–884. doi:10.1038/nature04723

7. Özcan U, Cao Q, Yilmaz E, et al. Endoplasmic Reticulum Stress Links Obesity, Insulin Action, and Type 2 Diabetes. Science (80-). 2004;306(5695):457 LP – 461. http://science.sciencemag.org/content/306/5695/457.abstract.

8. D’Angelo MA, Raices M, Panowski SH, Hetzer MW. Age-Dependent Deterioration of Nuclear Pore Complexes Causes a Loss of Nuclear Integrity in Postmitotic Cells. Cell. 2009;136(2):284–295. doi:10.1016/j.cell.2008.11.037

9. Lane RK, Hilsabeck T, Rea SL. The role of mitochondrial dysfunction in age-related diseases. Biochim Biophys Acta - Bioenerg. 2015;1847(11):1387–1400. doi:10.1016/j.bbabio.2015.05.021

10. Petersen KF, Dufour S, Befroy D, Garcia R, Shulman GI. Impaired Mitochondrial Activity in the Insulin-Resistant Offspring of Patients with Type 2 Diabetes. N Engl J Med. 2004;350(7):664–671. doi:10.1056/nejmoa031314

11. Mootha VK, Lindgren CM, Eriksson KF, et al. PGC-1α-responsive genes involved in oxidative phosphorylation are coordinately downregulated in human diabetes. Nat Genet. 2003;34(3):267–273. doi:10.1038/ng1180

12. Schapira AH V, Cooper JM, Dexter D, Clark JB, Jenner P, Marsden CD. Mitochondrial Complex I Deficiency in Parkinson’s Disease. J Neurochem. 1990;54(3):823–827. doi:10.1111/j.1471-4159.1990.tb02325.x

13. Bender A, Krishnan KJ, Morris CM, et al. High levels of mitochondrial DNA deletions in substantia nigra neurons in aging and Parkinson disease. Nat Genet. 2006;38(5):515–517. doi:10.1038/ng1769

14. Wanagat J, Cao Z, Pathare P, Aiken JM. Mitochondrial DNA deletion mutations colocalize with segmental electron transport system abnormalities, muscle fiber atrophy, fiber splitting, and oxidative damage in sarcopenia. FASEB J. 2001;15(2):322–332. doi:10.1096/fj.00-0320com

15. Ashar FN, Zhang Y, Longchamps RJ, et al. Association of mitochondrial DNA copy number with cardiovascular disease. JAMA Cardiol. 2017;2(11):1247–1255. doi:10.1001/jamacardio.2017.3683

16. Fleischman A, Makimura H, Stanley TL, et al. Skeletal muscle phosphocreatine recovery after submaximal exercise in children and young and middle-aged adults. J Clin Endocrinol Metab. 2010;95(9):69–74. doi:10.1210/jc.2010-0527

17. Fannin SW, Lesnefsky EJ, Slabe TJ, Hassan MO, Hoppel CL. Aging selectively decreases oxidative capacity in rat heart interfibrillar mitochondria. Arch Biochem Biophys. 1999;372(2):399–407. doi:10.1006/abbi.1999.1508

18. Trounce I, Byrne E, Marzuki S. Decline in Skeletal Muscle Mitochondrial Respiratory Chain Function: Possible Factor in Ageing. Lancet. 1989;333(8639):637–639. doi:10.1016/S0140-6736(89)92143-0

19. Kelley DE, He J, Menshikova E V., Ritov VB. Dysfunction of mitochondria in human skeletal muscle in type 2 diabetes. Diabetes. 2002;51(10):2944–2950. doi:10.2337/diabetes.51.10.2944

20. Patti ME, Butte AJ, Crunkhorn S, et al. Coordinated reduction of genes of oxidative metabolism in humans with insulin resistance and diabetes: Potential role of PGC1 and NRF1. Proc Natl Acad Sci U S A. 2003;100(14):8466–8471. doi:10.1073/pnas.1032913100

21. Stump CS, Short KR, Bigelow ML, Schimke JM, Nair KS. Effect of insulin on human skeletal muscle mitochondrial ATP production, protein synthesis, and mRNA transcripts. Proc Natl Acad Sci U S A. 2003;100(13):7996–8001. doi:10.1073/pnas.1332551100

22. Taylor RW, Barron MJ, Borthwick GM, et al. Mitochondrial DNA mutations in human colonic crypt stem cells Find the latest version : Mitochondrial DNA mutations in human colonic crypt stem cells. J Clin Invest. 2003;112(9):1351–1360. doi:10.1172/JCI200319435.Introduction

23. Frazier AE, Thorburn DR, Compton AG. Mitochondrial energy generation disorders: Genes, mechanisms, and clues to pathology. J Biol Chem. 2019;294(14):5386–5395. doi:10.1074/jbc.R117.809194

24. Curran JE, Johnson MP, Dyer TD, et al. Genetic determinants of mitochondrial content. Hum Mol Genet. 2007;16(12):1504–1514. doi:10.1093/hmg/ddm101

25. Xing J, Chen M, Wood CG, et al. Mitochondrial DNA content: Its genetic heritability and association with renal cell carcinoma. J Natl Cancer Inst. 2008;100(15):1104–1112. doi:10.1093/jnci/djn213

26. Kuan V, Denaxas S, Gonzalez-Izquierdo A, et al. A chronological map of 308 physical and mental health conditions from 4 million individuals in the English National Health Service. Lancet Digit Heal. 2019;1(2):e63–e77. doi:10.1016/s2589-7500(19)30012-3

27. Sudlow C, Gallacher J, Allen N, et al. UK Biobank: An Open Access Resource for Identifying the Causes of a Wide Range of Complex Diseases of Middle and Old Age. PLoS Med. 2015;12(3):1–10. doi:10.1371/journal.pmed.1001779

28. Teslovich TM, Musunuru K, Smith A V, et al. Biological, clinical and population relevance of 95 loci for blood lipids. Nature. 2010;466(7307):707–713. http://dx.doi.org/10.1038/nature09270.

29. Ehret GB, Munroe PB, Rice KM, et al. Genetic variants in novel pathways influence blood pressure and cardiovascular disease risk. Nature. 2011;478(7367):103–109. doi:10.1038/nature10405

30. Manning AK, Hivert M-F, Scott RA, et al. A genome-wide approach accounting for body mass index identifies genetic variants influencing fasting glycemic traits and insulin resistance. Nat Genet. 2012;44(6):659–669. http://dx.doi.org/10.1038/ng.2274.

31. Morris AP, Voight BF, Teslovich TM, et al. Large-scale association analysis provides insights into the genetic architecture and pathophysiology of type 2 diabetes. Nat Genet. 2012;44(9):981–990. doi:10.1038/ng.2383

32. Schunkert H, König IR, Kathiresan S, et al. Large-scale association analysis identifies 13 new susceptibility loci for coronary artery disease. Nat Genet. 2011;43(4):333–340. doi:10.1038/ng.784

33. Estrada K, Styrkarsdottir U, Evangelou E, et al. Genome-wide meta-analysis identifies 56 bone mineral density loci and reveals 14 loci associated with risk of fracture. Nat Genet. 2012;44(5):491–501. doi:10.1038/ng.2249

34. Christophersen IE, Rienstra M, Roselli C, et al. Large-scale analyses of common and rare variants identify 12 new loci associated with atrial fibrillation. Nat Genet. 2017;49(6):946–952. doi:10.1038/ng.3843

35. Pattaro C, Teumer A, Gorski M, et al. Genetic associations at 53 loci highlight cell types and biological pathways relevant for kidney function. Nat Commun. 2016;7:1–19. doi:10.1038/ncomms10023

36. Nalls MA, Blauwendraat C, Vallerga CL, et al. Identification of novel risk loci, causal insights, and heritable risk for Parkinson’s disease: a meta-analysis of genome-wide association studies. Lancet Neurol. 2019;18(12):1091–1102. doi:10.1016/S1474-4422(19)30320-5

37. Lambert JC, Ibrahim-Verbaas CA, Harold D, et al. Meta-analysis of 74,046 individuals identifies 11 new susceptibility loci for Alzheimer’s disease. Nat Genet. 2013;45(12):1452–1458. doi:10.1038/ng.2802

38. Finucane HK, Bulik-Sullivan B, Gusev A, et al. Partitioning heritability by functional annotation using genome-wide association summary statistics. Nat Genet. 2015;47(11):1228–1235. doi:10.1038/ng.3404

39. Bulik-Sullivan B, Finucane HK, Anttila V, et al. An atlas of genetic correlations across human diseases and traits. Nat Genet. 2015;47(11):1236–1241. doi:10.1038/ng.3406

40. Wasmer K, Eckardt L, Breithardt G. Predisposing factors for atrial fibrillation in the elderly. J Geriatr Cardiol. 2017;14(3):179–184. doi:10.11909/j.issn.1671-5411.2017.03.010

41. Calvo SE, Clauser KR, Mootha VK. MitoCarta2.0: An updated inventory of mammalian mitochondrial proteins. Nucleic Acids Res. 2016;44(D1):D1251–D1257. doi:10.1093/nar/gkv1003

42. MacArthur J, Bowler E, Cerezo M, et al. The new NHGRI-EBI Catalog of published genome-wide association studies (GWAS Catalog). Nucleic Acids Res. 2016;45(November 2016):gkw1133. doi:10.1093/nar/gkw1133

43. Finucane HK, Reshef YA, Anttila V, et al. Heritability enrichment of specifically expressed genes identifies disease-relevant tissues and cell types. Nat Genet. 2018;50(4):621–629. doi:10.1038/s41588-018-0081-4

44. de Leeuw CA, Mooij JM, Heskes T, Posthuma D. MAGMA: Generalized Gene-Set Analysis of GWAS Data. PLoS Comput Biol. 2015;11(4):1–19. doi:10.1371/journal.pcbi.1004219

45. Yamamoto K, Sakaue S, Matsuda K, et al. Genetic and phenotypic landscape of the mitochondrial genome in the Japanese population. Commun Biol. 2020;3(1):104. doi:10.1038/s42003-020-0812-9

46. Binder JX, Pletscher-Frankild S, Tsafou K, et al. COMPARTMENTS: Unification and visualization of protein subcellular localization evidence. Database. 2014;2014:1–9. doi:10.1093/database/bau012

47. Carbon S, Douglass E, Dunn N, et al. The Gene Ontology Resource: 20 years and still GOing strong. Nucleic Acids Res. 2019;47(D1):D330–D338. doi:10.1093/nar/gky1055

48. Ashburner M, Ball CA, Blake JA, et al. Gene Ontology: Tool for The Unification of Biology. Nat Genet. 2000;25(1):25–29. doi:10.1038/75556

49. Lambert SA, Jolma A, Campitelli LF, et al. The Human Transcription Factors. Cell. 2018;172(4):650–665. doi:10.1016/j.cell.2018.01.029

50. Kapopoulou A, Mathew L, Wong A, Trono D, Jensen JD. The evolution of gene expression and binding specificity of the largest transcription factor family in primates. Evolution (N Y). 2016;70(1):167–180. doi:10.1111/evo.12819

51. Timmers Prhj, Mounier N, Lall K, et al. Genomics of 1 million parent lifespans implicates novel pathways and common diseases and distinguishes survival chances. Elife. 2019;8:1–40. doi:10.7554/elife.39856

52. Zenin A, Tsepilov Y, Sharapov S, et al. Identification of 12 genetic loci associated with human healthspan. Commun Biol. 2019;2(1). doi:10.1038/s42003-019-0290-0

53. Jimenez-Sanchez G, Childs B, Valle D. Human Disease Genes. Nature. 2001;409:853–855.

54. Worman HJ, Courvalin JC. The nuclear lamina and inherited disease. Trends Cell Biol. 2002;12(12):591–598. doi:10.1016/S0962-8924(02)02401-7

55. Cleaver JE. It was a very good year for DNA repair. Cell. 1994;76(1):1–4. doi:10.1016/0092-8674(94)90165-1

56. Karczewski KJ, Francioli LC, Tiao G, et al. The mutational constraint spectrum quantified from variation in 141,456 humans. Nature. 2020;581(7809):434–443. doi:10.1038/s41586-020-2308-7

57. Colacurcio DJ, Nixon RA. Disorders of lysosomal acidification—The emerging role of v-ATPase in aging and neurodegenerative disease. Ageing Res Rev. 2016;32:75–88. doi:10.1016/j.arr.2016.05.004

58. Kanfi Y, Peshti V, Gil R, et al. SIRT6 protects against pathological damage caused by diet-induced obesity. Aging Cell. 2010;9(2):162–173. doi:10.1111/j.1474-9726.2009.00544.x

59. Blasco MA. Telomere length, stem cells and aging. Nat Chem Biol. 2007;3(10):640–649. doi:10.1038/nchembio.2007.38

60. Bhattarai KR, Chaudhary M, Kim H-R, Chae H-J. Endoplasmic Reticulum (ER) Stress Response Failure in Diseases. Trends Cell Biol. 2020;xx(xx):5–7. doi:10.1016/j.tcb.2020.05.004

61. De Leeuw CA, Neale BM, Heskes T, Posthuma D. The statistical properties of gene-set analysis. Nat Rev Genet. 2016;17(6):353–364. doi:10.1038/nrg.2016.29

62. Loh PR, Bhatia G, Gusev A, et al. Contrasting genetic architectures of schizophrenia and other complex diseases using fast variance-components analysis. Nat Genet. 2015;47(12):1385–1392. doi:10.1038/ng.3431

63. Billingsley KJ, Barbosa IA, Bandrés-Ciga S, et al. Mitochondria function associated genes contribute to Parkinson’s Disease risk and later age at onset. npj Park Dis. 2019;5(1). doi:10.1038/s41531-019-0080-x

64. Kraja AT, Liu C, Fetterman JL, et al. Associations of Mitochondrial and Nuclear Mitochondrial Variants and Genes with Seven Metabolic Traits. Am J Hum Genet. 2019;104(1):112–138. doi:10.1016/j.ajhg.2018.12.001

65. Xue A, Wu Y, Zhu Z, et al. Genome-wide association analyses identify 143 risk variants and putative regulatory mechanisms for type 2 diabetes. Nat Commun. 2018;9(1). doi:10.1038/s41467-018-04951-w

66. Jansen IE, Savage JE, Watanabe K, et al. Genome-wide meta-analysis identifies new loci and functional pathways influencing Alzheimer’s disease risk. Nat Genet. 2019;51(3):404–413. doi:10.1038/s41588-018-0311-9

67. Pardiñas AF, Holmans P, Pocklington AJ, et al. Common schizophrenia alleles are enriched in mutation-intolerant genes and in regions under strong background selection. Nat Genet. 2018;50(3):381–389. doi:10.1038/s41588-018-0059-2

68. Maurano MT, Humbert R, Rynes E, et al. Systematic localization of common disease-associated variation in regulatory DNA. Science (80-). 2012;337(6099):1190–1195. doi:10.1126/science.1222794

69. Raule N, Sevini F, Santoro A, Altilia S, Franceschi C. Association studies on human mitochondrial DNA: Methodological aspects and results in the most common age-related diseases. Mitochondrion. 2007;7(1-2):29–38. doi:10.1016/j.mito.2006.11.013

70. Yu X, Koczan D, Sulonen AM, et al. mtDNA nt13708A variant increases the risk of multiple sclerosis. PLoS One. 2008;3(2):1–7. doi:10.1371/journal.pone.0001530

71. Hudson G, Nalls M, Evans JR, et al. Two-stage association study and meta-analysis of mitochondrial DNA variants in Parkinson disease. Neurology. 2013;80(22):2042–2048. doi:10.1212/WNL.0b013e318294b434

72. Samuels DC, Carothers AD, Horton R, Chinnery PF. The power to detect disease associations with mitochondrial DNA haplogroups. Am J Hum Genet. 2006;78(4):713–720. doi:10.1086/502682

73. Biffi A, Anderson CD, Nalls MA, et al. Principal-Component Analysis for Assessment of Population Stratification in Mitochondrial Medical Genetics. Am J Hum Genet. 2010;86(6):904–917. doi:10.1016/j.ajhg.2010.05.005

74. Segrè A V, Consortium D, investigators M, et al. Common Inherited Variation in Mitochondrial Genes Is Not Enriched for Associations with Type 2 Diabetes or Related Glycemic Traits. PLOS Genet. 2010;6(8):e1001058. https://doi.org/10.1371/journal.pgen.1001058.

75. Saxena R, De Bakker PIW, Singer K, et al. Comprehensive association testing of common mitochondrial DNA variation in metabolic disease. Am J Hum Genet. 2006;79(1):54–61. doi:10.1086/504926

76. Hudson G, Gomez-Duran A, Wilson IJ, Chinnery PF. Recent Mitochondrial DNA Mutations Increase the Risk of Developing Common Late-Onset Human Diseases. PLoS Genet. 2014;10(5). doi:10.1371/journal.pgen.1004369

77. Hudson G, Panoutsopoulou K, Wilson I, et al. No evidence of an association between mitochondrial DNA variants and osteoarthritis in 7393 cases and 5122 controls. Ann Rheum Dis. 2013;72(1):136–139. doi:10.1136/annrheumdis-2012-201932

78. Seidman JG, Seidman C. Transcription factor haploinsufficiency: When half a loaf is not enough. J Clin Invest. 2002;109(4):451–455. doi:10.1172/JCI0215043

79. Lee R Van Der, Correard S, Wasserman WW. Deregulated Regulators : Disease-Causing cis Variants in Transcription Factor Genes. Trends Genet. 2020;xx(xx). doi:10.1016/j.tig.2020.04.006

80. Wright S. Physiological and Evolutionary Theories of Dominance. Am Nat. 1934;68(714):24.

81. Kacser H, Burns JA. The molecular basis of dominance. Genetics. 1981;97(3-4):639–666.

82. Chance B, Williams GR. RESPIRATORY ENZYMES IN OXIDATIVE PHOSPHORYLATION: III. THE STEADY STATE. J Biol Chem. 1955;217(1):409–428. http://www.jbc.org/content/217/1/409.short.

83. Vafai SB, Mootha VK. Mitochondrial disorders as windows into an ancient organelle. Nature. 2012;491(7424):374–383. doi:10.1038/nature11707

84. Balaban RS, Kantor HL, Katz LA, Briggs RW. Relation between work and phosphate metabolite in the in vivo paced mammalian heart. Science (80-). 1986;232(4754):1121–1123. doi:10.1126/science.3704638

85. To TL, Cuadros AM, Shah H, et al. A Compendium of Genetic Modifiers of Mitochondrial Dysfunction Reveals Intra-organelle Buffering. Cell. 2019;179(5):1222-1238.e17. doi:10.1016/j.cell.2019.10.032

86. Golan D, Lander ES, Rosset S. Measuring missing heritability: Inferring the contribution of common variants. Proc Natl Acad Sci U S A. 2014;111(49):E5272–E5281. doi:10.1073/pnas.1419064111

87. Polderman TJC, Benyamin B, De Leeuw CA, et al. Meta-analysis of the heritability of human traits based on fifty years of twin studies. Nat Genet. 2015;47(7):702–709. doi:10.1038/ng.3285

88. Fuchsberger C, Flannick J, Teslovich TM, et al. The genetic architecture of type 2 diabetes. Nature. 2016;536(7614):41–47. doi:10.1038/nature18642

89. Hill WG, Goddard ME, Visscher PM. Data and theory point to mainly additive genetic variance for complex traits. PLoS Genet. 2008;4(2). doi:10.1371/journal.pgen.1000008

90. Zhu Z, Bakshi A, Vinkhuyzen AAE, et al. Dominance genetic variation contributes little to the missing heritability for human complex traits. Am J Hum Genet. 2015;96(3):377–385. doi:10.1016/j.ajhg.2015.01.001

91. Rand DM, Mossman JA. Mitonuclear conflict and cooperation govern the integration of genotypes, phenotypes and environments. Philos Trans R Soc B Biol Sci.2020;375(1790). doi:10.1098/rstb.2019.0188

92. Sackton TB, Hartl DL. Genotypic Context and Epistasis in Individuals and Populations. Cell. 2016;166(2):279–287. doi:10.1016/j.cell.2016.06.047

93. Hemani G, Shakhbazov K, Westra HJ, et al. Detection and replication of epistasis influencing transcription in humans. Nature. 2014;508(7495):249–253. doi:10.1038/nature13005

94. Solenski NJ, DiPierro CG, Trimmer PA, Kwan AL, Helms GA. Ultrastructural changes of neuronal mitochondria after transient and permanent cerebral ischemia. Stroke. 2002;33(3):816–824. doi:10.1161/hs0302.104541

95. Flameng W, Andres J, Ferdinande P, Mattheussen M, Van Belle H. Mitochondrial function in myocardial stunning. J Mol Cell Cardiol. 1991;23(1):1–11. doi:10.1016/0022-2828(91)90034-J

96. Weinbrenner C, Liu GS, Downey JM, Cohen M V. Cyclosporine a limits myocardial infarct size even when administered after onset of ischemia. Cardiovasc Res. 1998;38(3):676–684. doi:10.1016/S0008-6363(98)00064-9

97. Kubben N, Misteli T. Shared molecular and cellular mechanisms of premature ageing and ageing-associated diseases. Nat Rev Mol Cell Biol. 2017;18(10):595–609. doi:10.1038/nrm.2017.68

98. Garcia CK, Wright WE, Shay JW. Human diseases of telomerase dysfunction: Insights into tissue aging. Nucleic Acids Res. 2007;35(22):7406–7416. doi:10.1093/nar/gkm644

99. Han S, Brunet A. Histone methylation makes its mark on longevity. Trends Cell Biol. 2012;22(1):42–49. doi:10.1016/j.tcb.2011.11.001

100. Bahar R, Hartmann CH, Rodriguez KA, et al. Increased cell-to-cell variation in gene expression in ageing mouse heart. Nature. 2006;441(7096):1011–1014. doi:10.1038/nature04844

101. Kanfi Y, Naiman S, Amir G, et al. The sirtuin SIRT6 regulates lifespan in male mice. Nature. 2012;483(7388):218–221. doi:10.1038/nature10815

102. Karczewski KJ, Dudley JT, Kukurba KR, et al. Systematic functional regulatory assessment of disease-associated variants. Proc Natl Acad Sci U S A. 2013;110(23):9607–9612. doi:10.1073/pnas.1219099110

103. Yanai I, Benjamin H, Shmoish M, et al. Genome-wide midrange transcription profiles reveal expression level relationships in human tissue specification. Bioinformatics. 2005;21(5):650–659. doi:10.1093/bioinformatics/bti042

104. Melé M, Ferreira PG, Reverter F, et al. The human transcriptome across tissues and inidividuals. Science (80-). 2015;348(6235):660–665. http://dx.doi.org/10.1126/science.aaa0355.

105. Morris JA, Randall JC, Maller JB, Barrett JC. Evoker: A visualization tool for genotype intensity data. Bioinformatics. 2010;26(14):1786–1787. doi:10.1093/bioinformatics/btq280

106. Yates AD, Achuthan P, Akanni W, et al. Ensembl 2020. Nucleic Acids Res. 2020;48(D1):D682–D688. doi:10.1093/nar/gkz966

107. Litman T, Stein WD. Obtaining estimates for the ages of all the protein-coding genes and most of the ontology-identified noncoding genes of the human genome, assigned to 19 phylostrata. Semin Oncol. 2019;46(1):3–9. doi:10.1053/j.seminoncol.2018.11.002

