## Supplemental figures, Table S1, Supplementary text for "Human genetic analyses of organelles highlight the nucleus in age-related trait heritability"

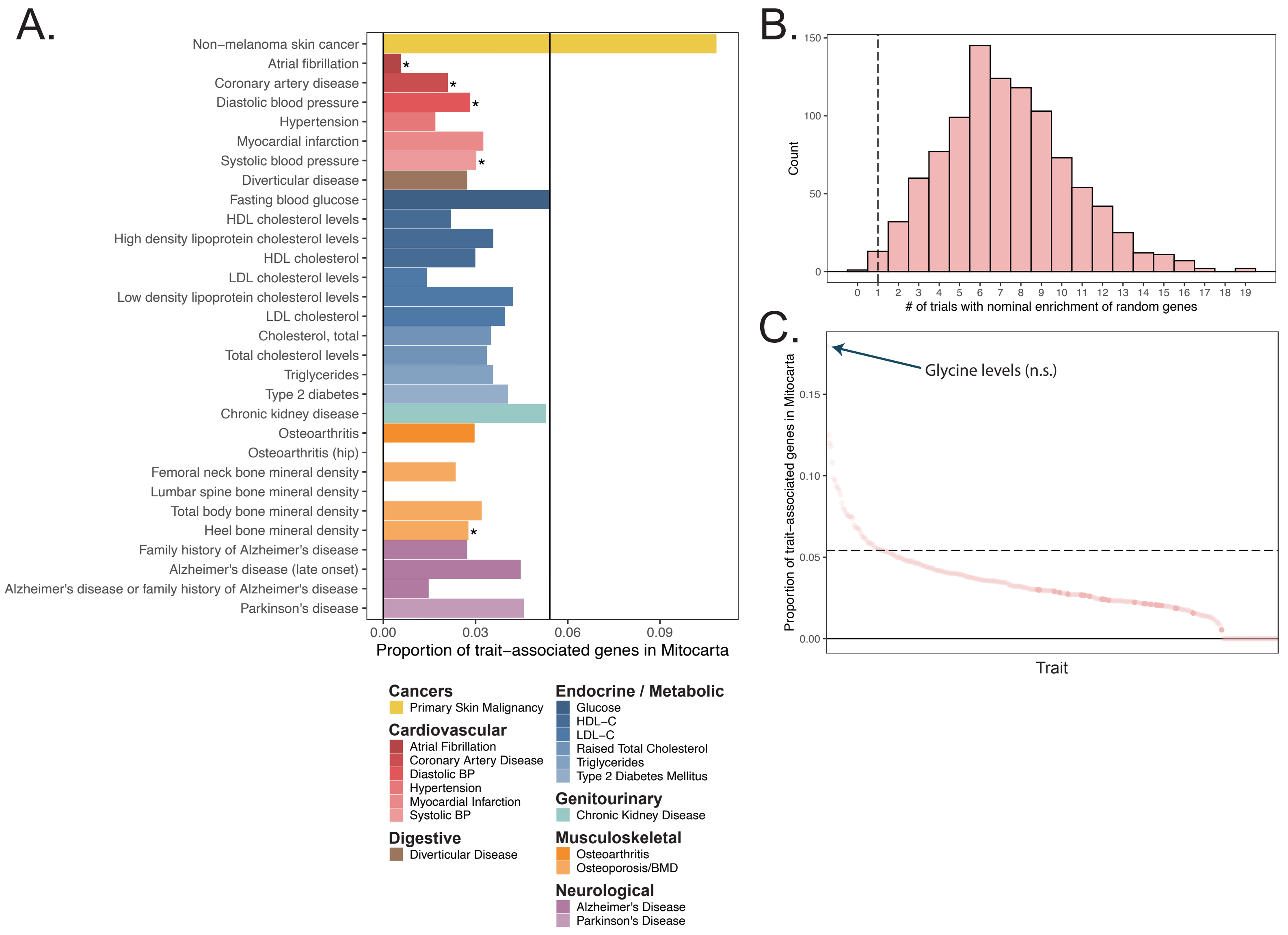


**Figure 2-S1.** Enrichment tests in mitochondria-localizing genes in the GWAS Catalog. **A.** The proportion of annotated lead SNP-associated genes for manually curated phenotypes from the GWAS Catalog that correspond to age-related traits of interest (colors) that overlap the set of mitochondria-localizing genes. Black line represents the proportion of all genes in the genome that are mitochondria-localizing, with bars above the black line representing nominal enrichment and below representing nominal depletion. * represents traits showing significant depletion or enrichment via two-sided Fisher's exact test at BH FDR q-value < 0.1. **B.** Empirical null distribution of the test statistic of the number of traits with nominal enrichment computed via 1,000 randomly sampled subsets of the set of all protein-coding genes. Dotted line represents the true observed value for mitochondria-localizing genes. **C.** Rank plot of the proportion of trait-associated genes overlapping with mitochondria-localizing genes (the same values as represented on the x-axis of panel **A** across all GWAS catalog phenotypes, only showing those with at least 30 trait-associated genes. Dark points indicate traits with a significant depletion or enrichment via two-sided Fisher's exact test at BH FDR q-value < 0.1. Dotted line represents the proportion of all genes in the genome that are mitochondria-localizing, with traits above the black line representing nominal enrichment and below representing nominal depletion. No traits show significant enrichment.

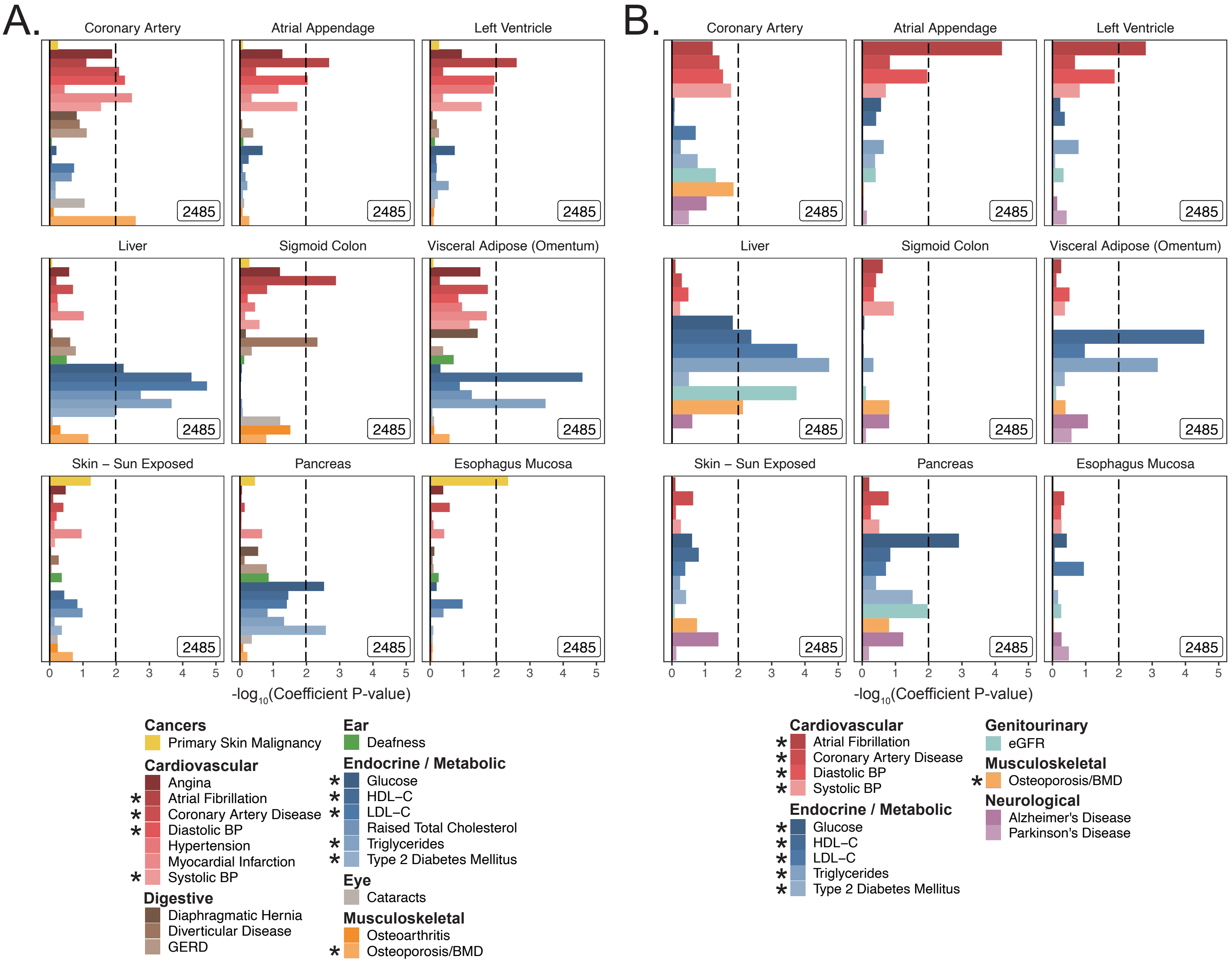


**Figure 2-S2.** Tissue enrichments in **A.** UK Biobank and **B.** meta-analyses using S-LDSC on top of the baseline model. Gene-sets of the top 10% tissue-specific genes for each tested tissue were obtained using GTEx v7 as described previously (Finucane et al. 2018), with tissues for analysis selected manually based on the set of tested traits. * represents traits for which sufficiently well powered cohorts from both UKB and meta-analyses were available. Dashed line represents $-\log_{10}$ P-value cutoff for Benjamini-Hochberg FDR < 10%.

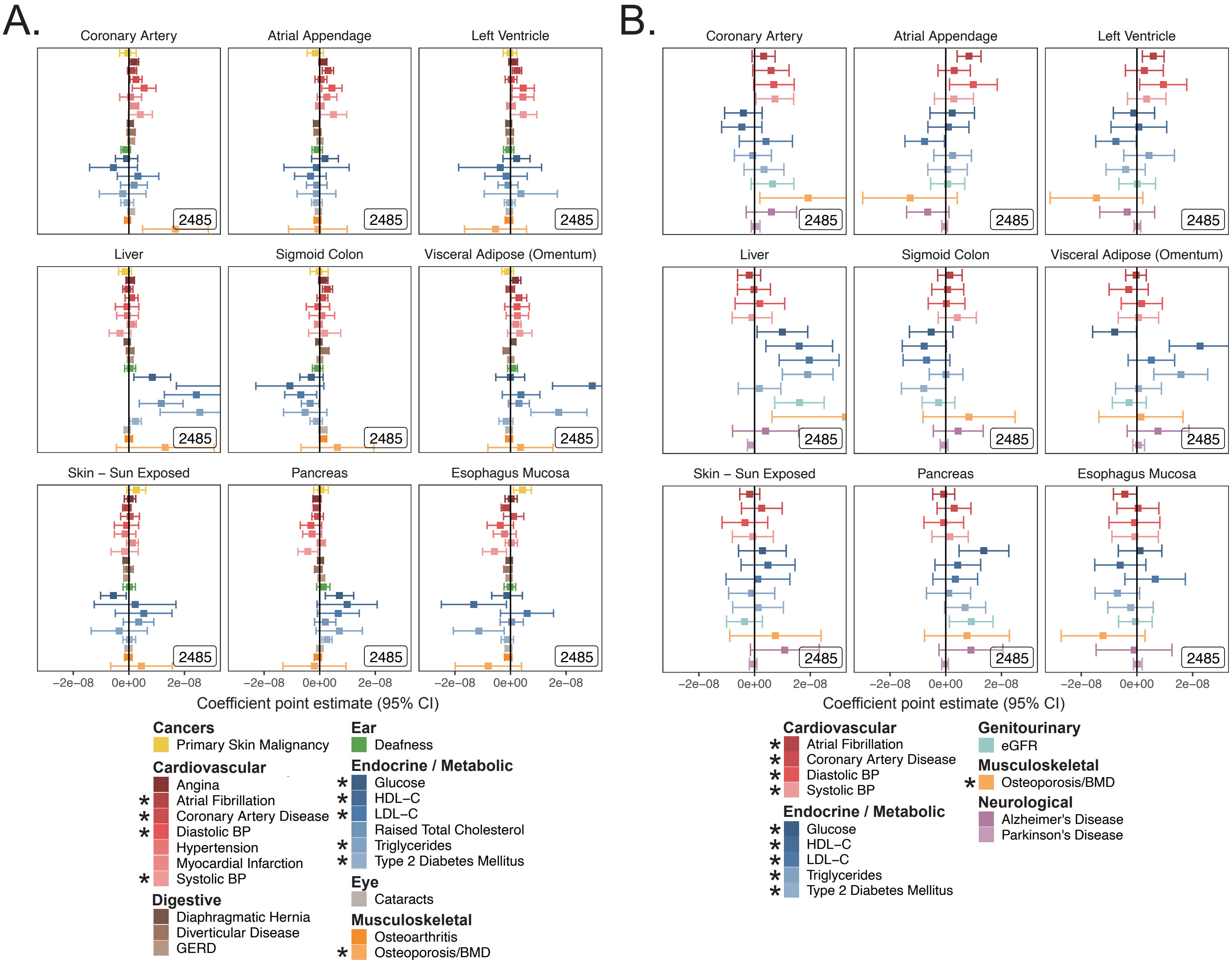


**Figure 2-S3.** S-LDSC enrichment coefficients from tissue analysis. **A.** Coefficient point estimates and corresponding 95% CI for tissue-expressed gene-sets in UKB with p-values shown in **Figure 2-S2. B.** Coefficient point estimates and 95% CI for tissue-expressed gene-sets tested in meta-analyses with p-values shown in **Figure 2-S2.** Inset numbers represent gene-set sizes. * represents traits for which sufficiently well powered cohorts from both UKB and meta-analyses were available. All analyses were performed atop the baseline model.

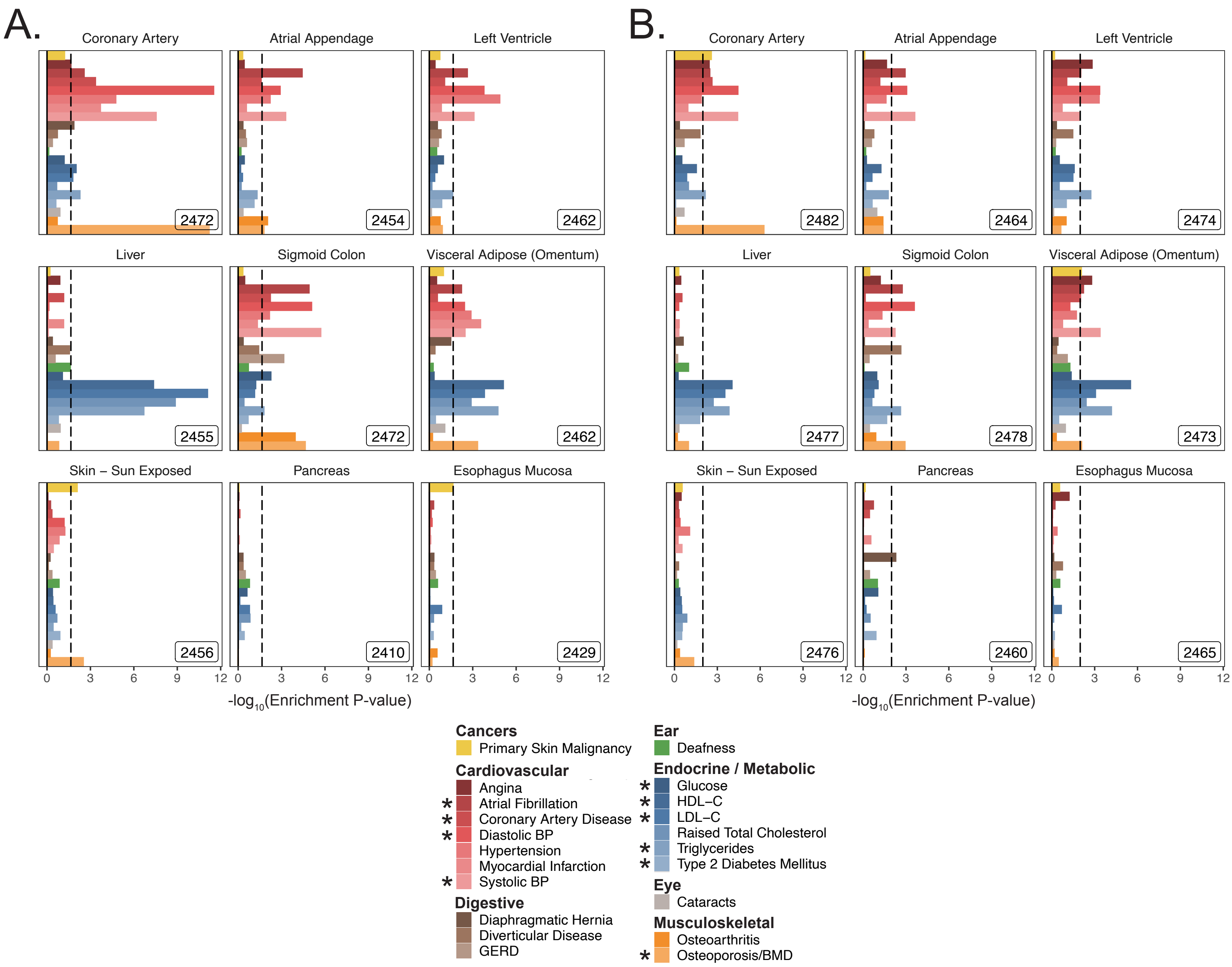


**Figure 2-S4.** MAGMA enrichments for tissues in UKB. Enrichment P-values for tissues with a **A.** 5kb up and 1.5kb down window and **B.** 100kb symmetric window. These gene-sets are analogous to those shown tested with S-LDSC in **Figure 2-S2** and **2-S3**. Inset numbers represent gene-set sizes. Dashed line represents –log10 P-value cutoff for Benjamini-Hochberg FDR < 10%. * represents traits for which sufficiently well powered cohorts from both UKB and meta-analyses were available.

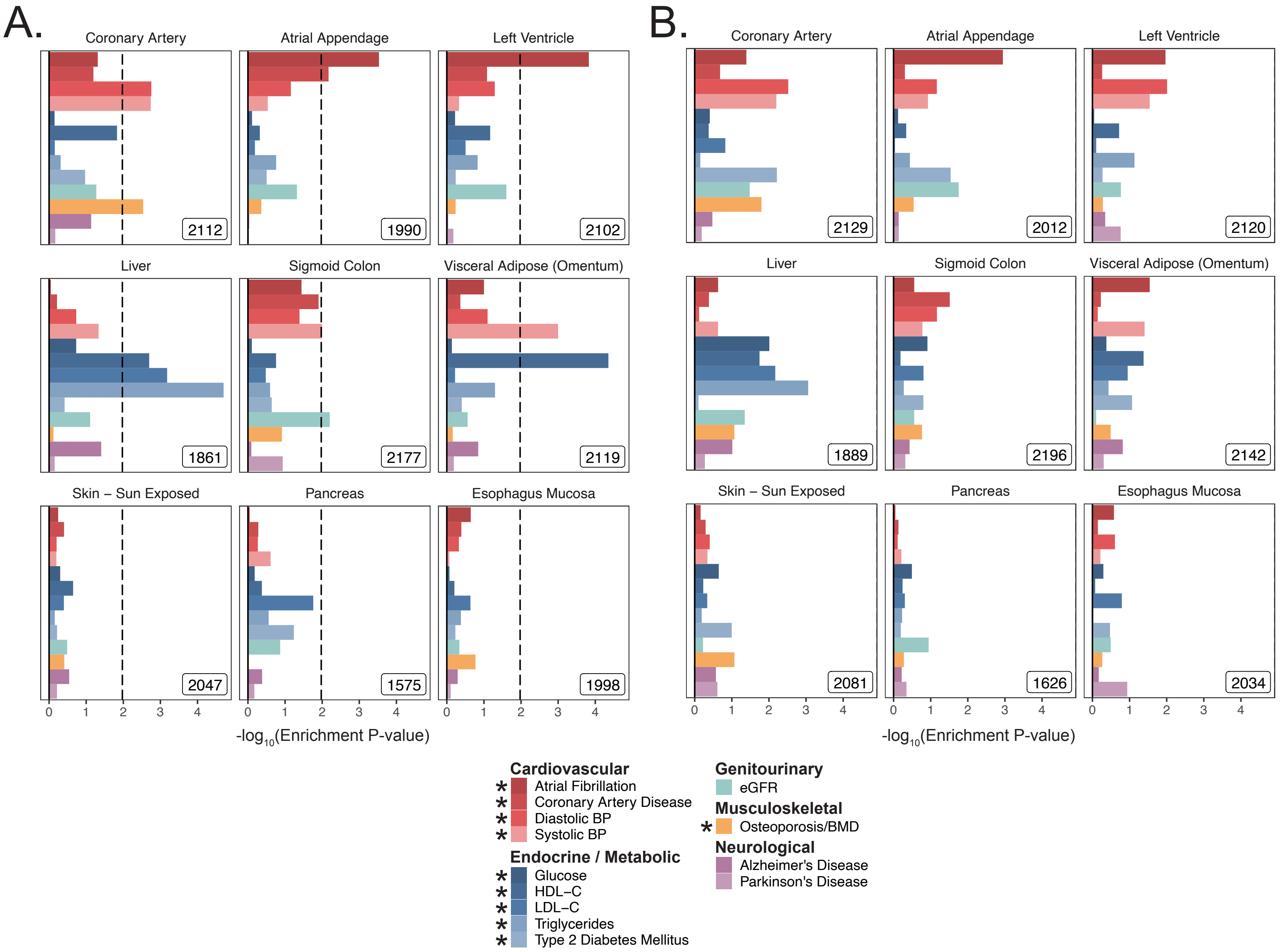


**Figure 2-S5.** MAGMA enrichments for tissues in meta-analyses. Enrichment P-values for tissues with a **A.** 5kb up and 1.5kb down window and **B.** 100kb symmetric window. These gene-sets are analogous to those shown tested with S-LDSC in **Figure 2-S2** and **2-S3** and with MAGMA in **Figure 2-S4**. Dashed line represents –log10 P-value cutoff for Benjamini-Hochberg FDR < 10%. No p-values crossed the FDR threshold for the 100kb window, and as such the BH P-value cutoff cannot be displayed. * represents traits for which sufficiently well powered cohorts from both UKB and meta-analyses were available.

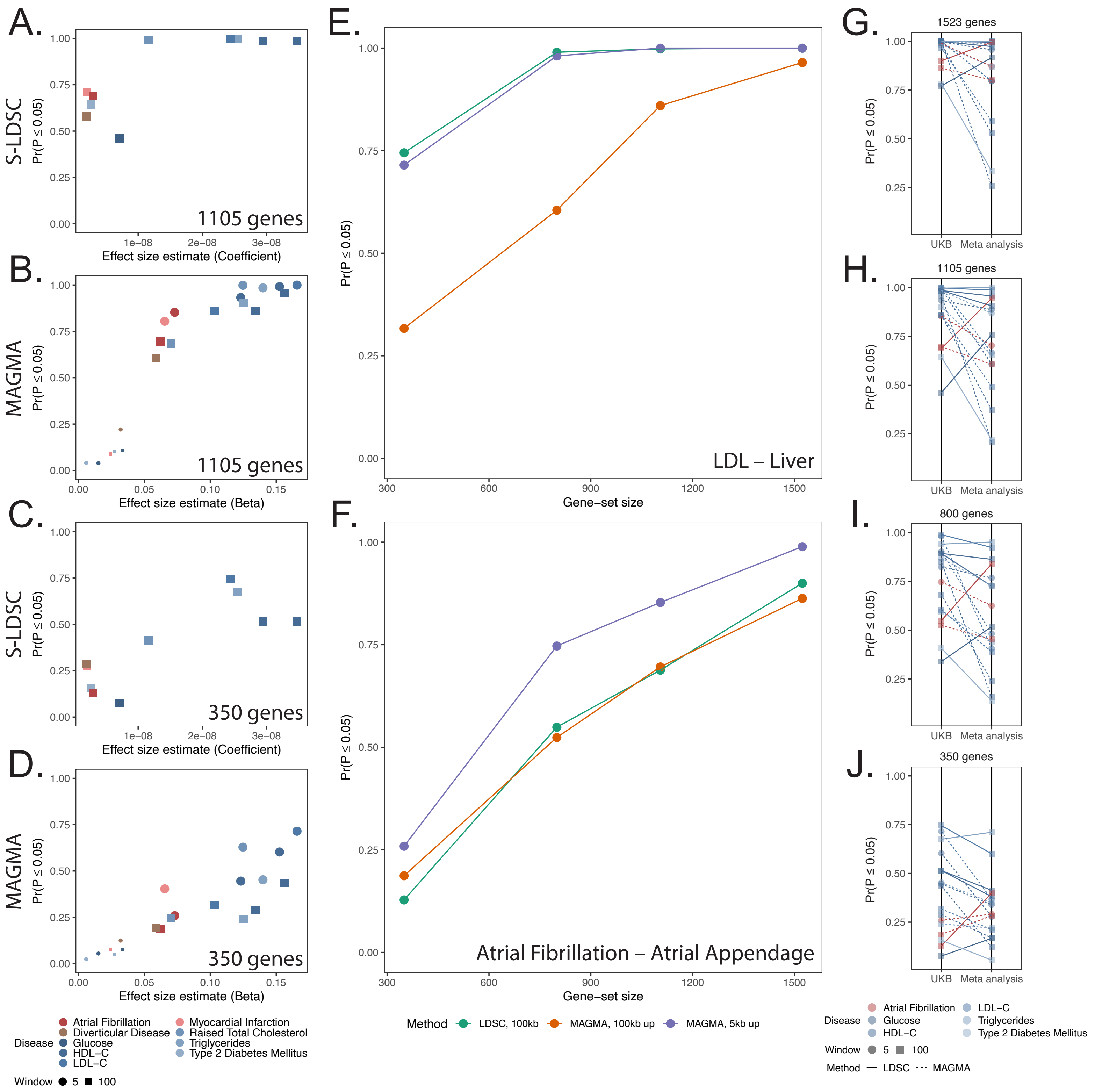


**Figure 2-S6.** Enrichment power of S-LDSC and MAGMA across effect sizes and gene-set sizes. Using biologically plausible trait-tissue associations originally computed at gene-set sizes of ~2500 genes (Atrial Fibrillation – Atrial appendage; Diverticular Disease – Sigmoid colon; Glucose, T2D – Pancreas; HDL, LDL, Cholesterol, TG – Liver, HDL – Visceral adipose; Myocardial Infarction – Coronary artery), power was computed as a probability of rejection at $P \leq 0.05$ across 1000 randomly sampled subsets (of size 1520, 1105, 800, and 350 genes) of the original tissue expression gene-set. **A.** Relationship between S-LDSC coefficient magnitude and power for 1105 genes. **B.** Relationship between MAGMA effect size magnitude and power for 1105 genes. **C.** Relationship between S-LDSC coefficient magnitude and power for 350 genes. **D.** Relationship between MAGMA effect size magnitude and power for 350 genes. Shape represents window size, color represents disease, and small points represent trait-tissue pairs that were not detected as enriched at the full gene-set. **E.** Power as a function of gene-set size for MAGMA and S-LDSC for a very high effect size trait-tissue pair: LDL levels and liver. **F.** Power as a function of gene-set size for MAGMA and S-LDSC for a low effect size trait-tissue pair: Atrial fibrillation and atrial appendage. UKB vs. meta-analysis power comparisons for **G.** 1523 genes, **H.** 1105 genes, **I.** 800 genes, and **J.** 350 genes.

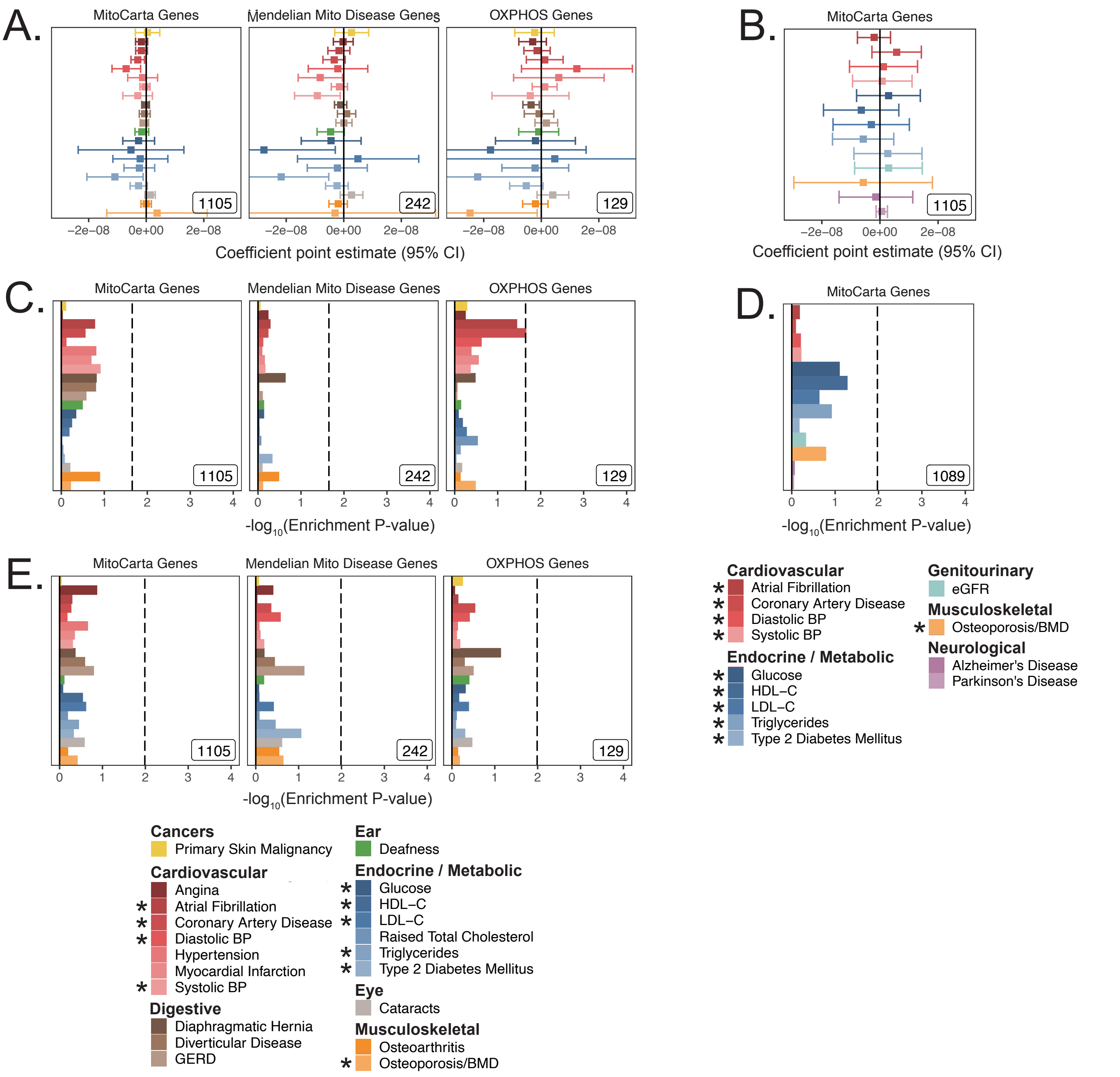


**Figure 2-S7.** Analysis of mitochondria-localizing gene enrichments using alternative methods and cohorts. **A.** Coefficient point estimates and corresponding 95% CI for mitochondria-localizing gene-sets in UKB with p-values shown in **Figure 2B**. **B.** Coefficient point estimates and 95% CI for mitochondria-localizing gene-set tested for replication in meta-analyses with p-values shown in **Figure 2C**. MAGMA enrichments for mitochondria-localizing genes with 5kb up and 1.5kb down gene window in **C.** UKB and **D.** meta-analyses. **E.** MAGMA enrichments for mitochondria-localizing genes in UKB with a 100kb symmetric gene window. Dotted line represents BH FDR 0.1 threshold. * represents traits for which sufficiently well powered cohorts from both UKB and meta-analyses were available. Inset numbers represent gene-set sizes.

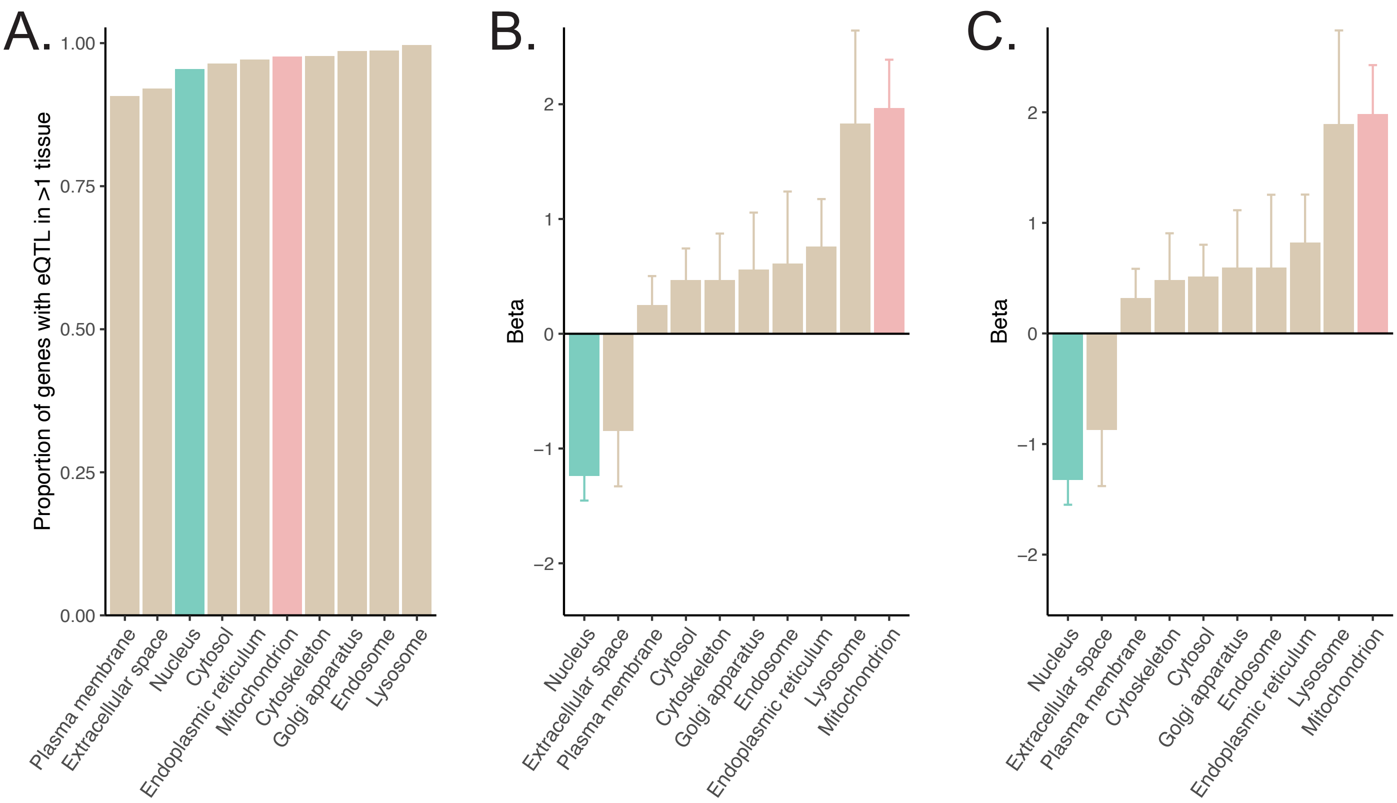


**Figure 2-S8.** Assessment of cis-eQTLs among mitochondria-localizing genes. **A.** Percentage of genes localizing to each organelle that have an observed cis-eQTL in at least one GTEx tissue. **B.** Coefficient effect size estimates for the regression of the number of distinct tissues with detected cis-eQTL for a given gene onto an indicator for organelle membership (**Methods**). **C.** Replication of **B** with alternate selection of distinct tissues for robustness (**Supplementary note**). Error bars represent 95% CI.

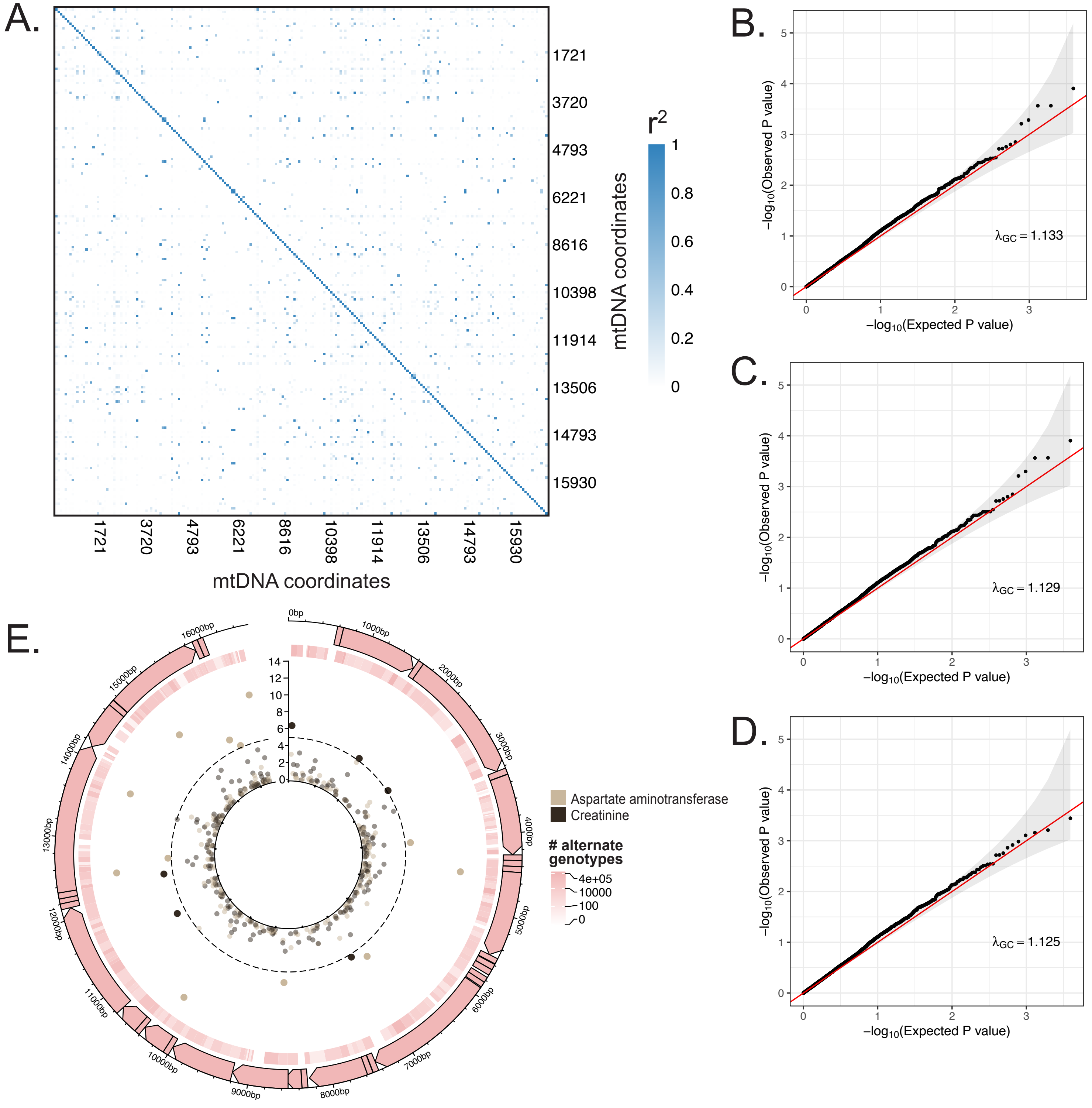


**Figure 2-S9.** Landscape of mtDNA variants and their associations with age-related disease in UKB. **A.** Within-mtDNA pair-wise correlation coefficients across all 213 tested variants with expected case count of alternative genotype individuals > 20. Axis labels represent mtDNA coordinates, color represents $r^2$. **B.** Quantile-quantile plot of p-values for all tested variant-trait pairs (21 traits x ~213 variants = ~4473 tests) using linear regression for all traits while controlling for baseline characteristics (first 20 PCs of nuclear genotype matrix, age, sex, age*sex, $age^2$ *sex) as well as array type. **C.** Quantile-quantile plot of all tested variant-trait pairs for all traits using linear regression, controlling only for baseline characteristics. **D.** Quantile-quantile plot of all tested variant-trait pairs using linear regression for continuous traits and logistic regression for binary traits, controlling only for baseline characteristics. Red line represents expected null p-values following the uniform distribution and shaded ribbon represents 95% CI. **E.** Visualization of mtDNA variants and associations with serum creatinine and aspartate aminotransferase levels. The outer-most track represents the genetic architecture of the circular mtDNA. The heatmap track represents the number of individuals with alternate genotype on log scale. The inner track represents mitochondrial genome-wide association p-values, with radial angle corresponding to position on the mtDNA and magnitude representing $-\log_{10}$ P-value. Dotted line represents Bonferroni cutoff for all tested trait-variant pairs, using the same threshold as used in **Figure 2D**.

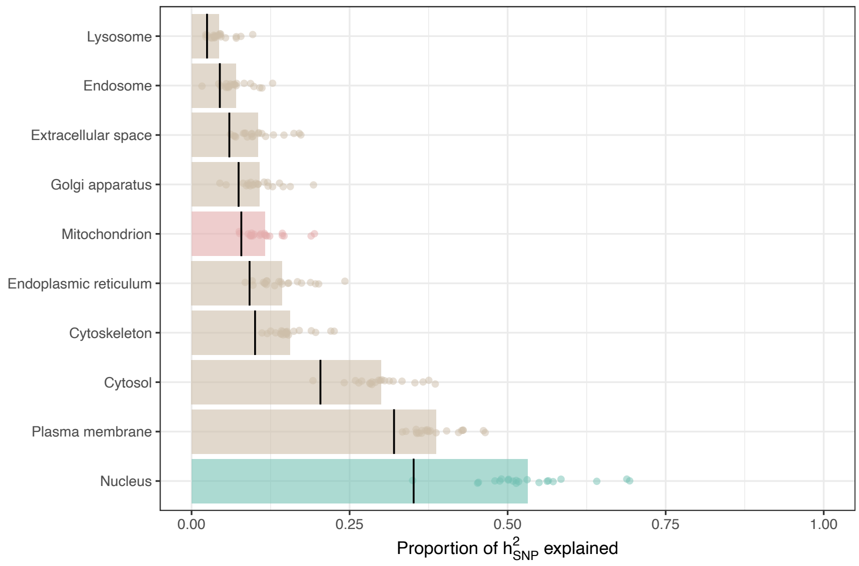


**Figure 3-S1.** Proportions of heritability explained by annotations indicating organellar localization as obtained via S-LDSC. All categories are identical to those analyzed in **Figure 3A**, however are obtained without controlling for the contributions to heritability from other categories in the baseline model. Each dot represents one of the 21 UKB age-related traits tested, with bar size determined by mean proportion of heritability explained. Black lines indicate the proportion of tested SNPs contained in each annotation.

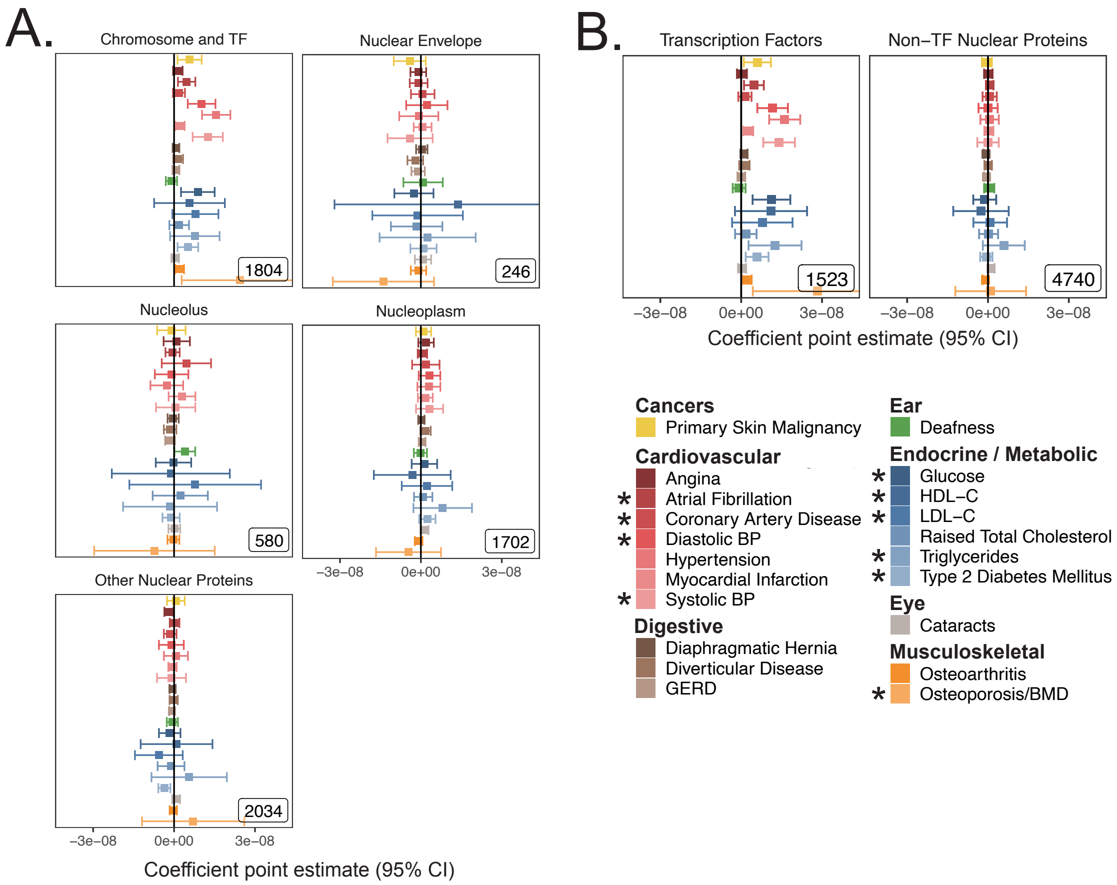


**Figure 3-S2.** S-LDSC coefficients from sub-nuclear compartment analysis. **A.** S-LDSC coefficient point estimates and 95% CI for spatially determined sub-categories of the nucleus as in **Figure 3C**. **B.** S-LDSC coefficient point estimates and 95% CI for partitions of the transcription factors and the rest of the nuclear proteome as in **Figure 3D**. Inset numbers represent gene-set size. * represents traits for which sufficiently well powered cohorts from both UKB and meta-analyses were available.

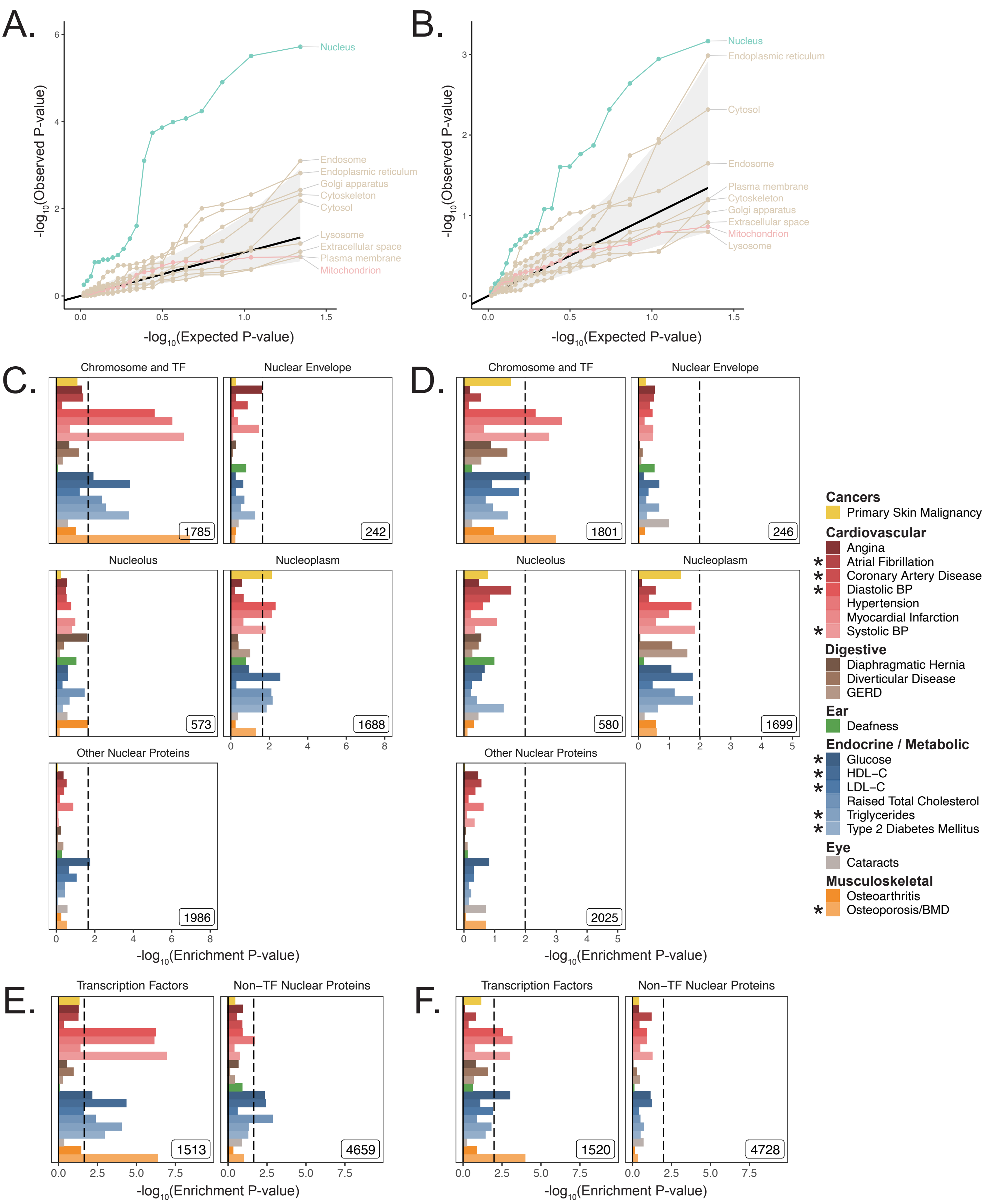


**Figure 3-S3.** Organelle analysis results in UKB using MAGMA. Quantile-quantile plot of one-sided enrichment P-values across UKB age-related traits using the same organelle gene-sets as in **Figure 3A**, with a **A.** 5kb up, 1.5kb down gene window and **B.** 100kb symmetric gene window. The central black line represents expected null p-values following the uniform distribution and the shaded ribbon represents 95% CI. **C.** Enrichment of spatially distinct disjoint subsets of the nuclear proteome as tested with S-LDSC in **Figure 3C**, using MAGMA with a 5kb up, 1.5kb down gene window and **D.** with a 100kb symmetric gene window. **E.** Enrichment of the transcription factors and the rest of the nuclear proteome as in **Figure 3D** using MAGMA with a 5kb up, 1.5kb down gene window and **F.** with a 100kb symmetric gene window. Inset numbers represent gene-set sizes, black lines represent cutoff at BH FDR < 10%.

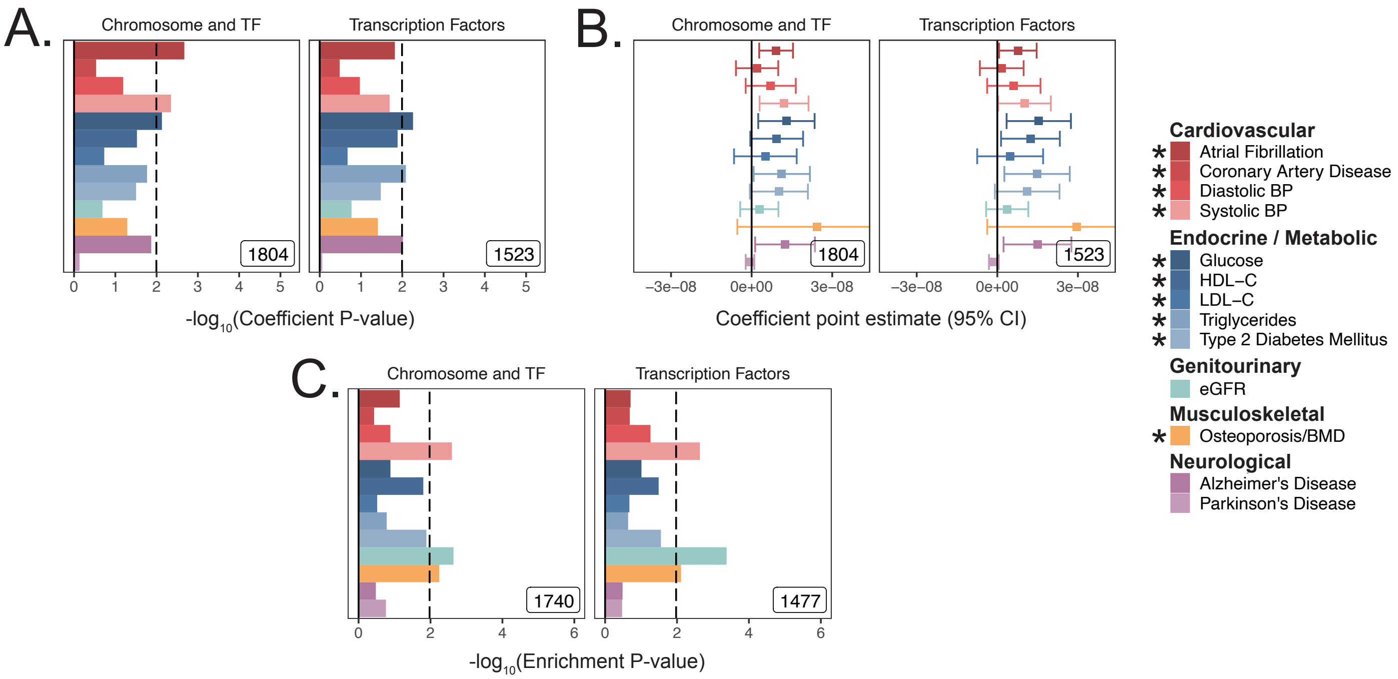


**Figure 3-S4.** Replication of transcription factor enrichment in meta-analyses. **A.** Coefficient P-values atop the baseline using S-LDSC model. **B.** Coefficient point-estimates corresponding to panel **A**. Error bars represent 95% CI. **C.** Enrichment P-values in meta-analyses using MAGMA with a 5kb up, 1.5kb down window. The gene-sets used for this analysis are identical to those tested in UKB in **Figure 3D**. * represents traits for which sufficiently well powered cohorts from both UKB and meta-analyses were available. Black lines represent cutoff at BH FDR < 10%. Inset numbers represent gene-set sizes.

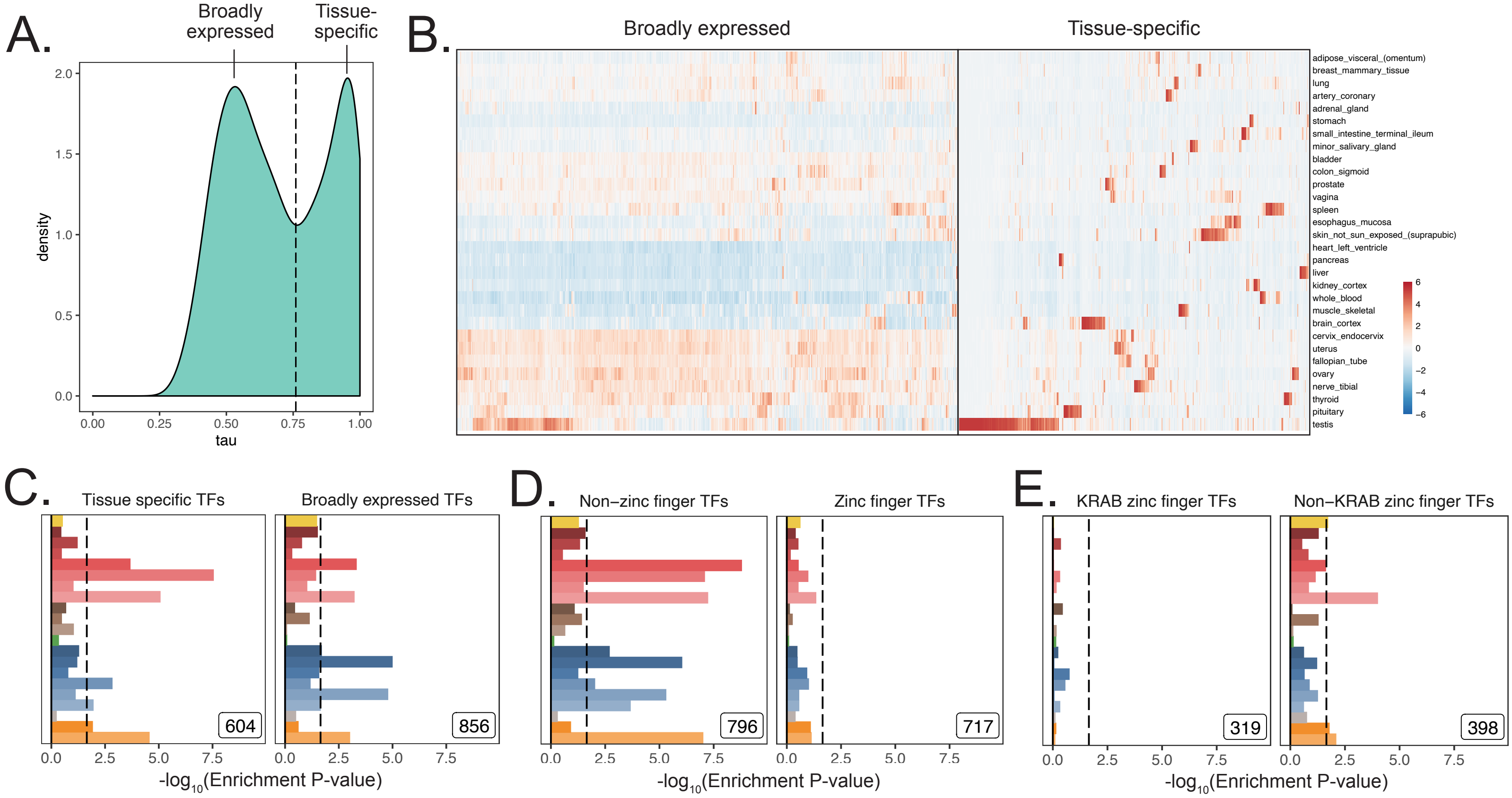


F.

| Ancient | Intermediate-age | Recently-evolved |
|---|---|---|
| Bilateria* | Amniota** | Primata (recent)** |
| Eumetazoa* | Tetrapoda** | Primata (old)** |
| Metazoa* | Euteleostomi** | Euarchonotoglires** |
| Holozoa* | Craniata** | Boreoeutheria** |
| Opisthokonta* | Olfactores* | Eutheria** |
| Eurkaryota* | Chordata* | Mammalia (recent)** |
| Last known common ancestor* | Deuterostomia* | Mammalia (old)** |

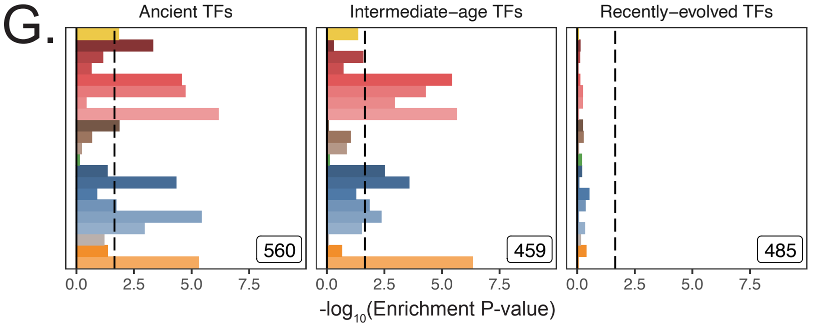


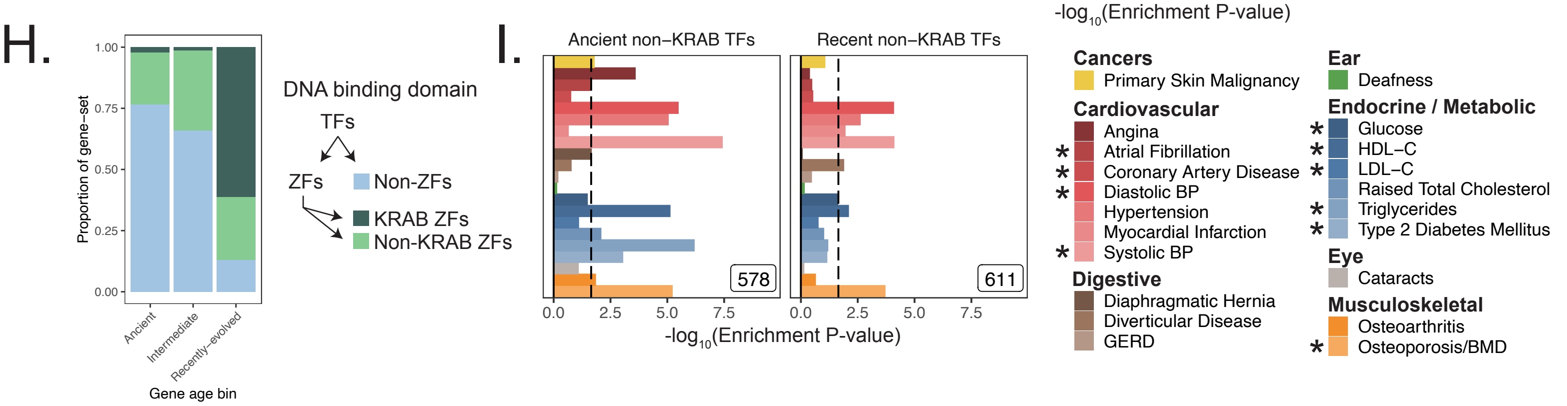


**Figure 3-S5.** MAGMA enrichments of functional subsets of transcription factors in UKB. **A.** Distribution of Tau statistic across genes, as computed in GTEx V7 (**Methods**). Black line indicates location of cutoff between tissue-specific genes (high Tau) and broadly expressed genes (low Tau). **B.** GTEx v7 medians per tissue across tissues used to construct broadly expressed and tissue-specific transcription factor gene-sets. Rows were ordered by unbiased hierarchical clustering via complete linkage across all genes; columns were ordered using hierarchical clustering within broadly-expressed and tissue-specific genes. Expression values are TPMs normalized across tissues. **C.** Enrichment analysis of TFs subdivided by breadth of expression as defined in **A.** **D.** Enrichment analysis of TFs subdivided by the presence of a zinc finger domain. **E.** Enrichment analysis of zinc finger TFs, subdivided by presence of KRAB domain. **F.** Phylostrata comprising each tested gene age bins used in **G**, **H**, **I**. Each gene was assigned a phylostratum based on its oldest ortholog by Litman & Stein, 2018. * is phylostratum that was placed into Ancient non-KRAB TFs; ** is a phylostratum placed into Recent non-KRAB TFs. **G.** Enrichment results dividing TFs into terciles based on evolutionary gene age. **H.** TF DNA binding domains represented in each gene age bin tested in panel **G**. **I.** Enrichment results subdividing TFs by age after removing KRAB domain zinc finger TFs. Inset numbers represent gene-set size, black lines for enrichment plots represent cutoff at BH FDR < 10%. All analyses were conducted with 5kb up, 1.5kb down windows.

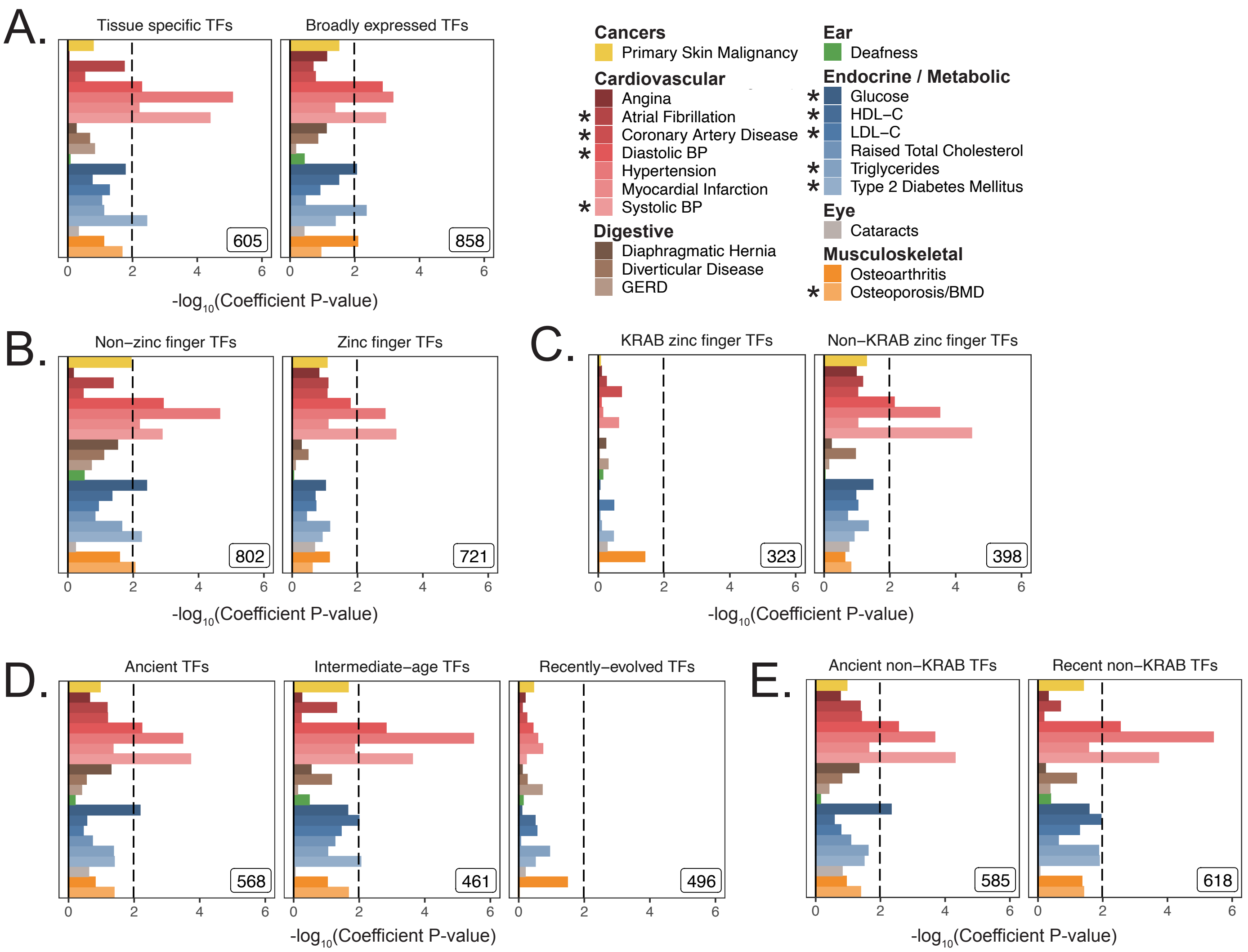


**Figure 3-S6.** Replication of enrichment analysis in functional subsets of transcription factors using S-LDSC. **A.** Enrichment analysis of TFs, subdivided by breadth of tissue expression in GTEx v7 (**Figure 3-S5A**, **3-S5B**). **B.** Enrichment analysis of TFs subdivided by the presence of a zinc finger domain. **C.** Enrichment analysis of zinc finger TFs, subdivided by presence of KRAB domain. **D.** Heritability enrichment results dividing TFs into terciles based on evolutionary gene age (see **Figure 3-S5F**). **E.** Enrichment analysis subdividing TFs into old and new after removing KRAB domain zinc finger TFs. Inset numbers represent gene-set size, black lines represent cutoff at BH FDR < 10%.

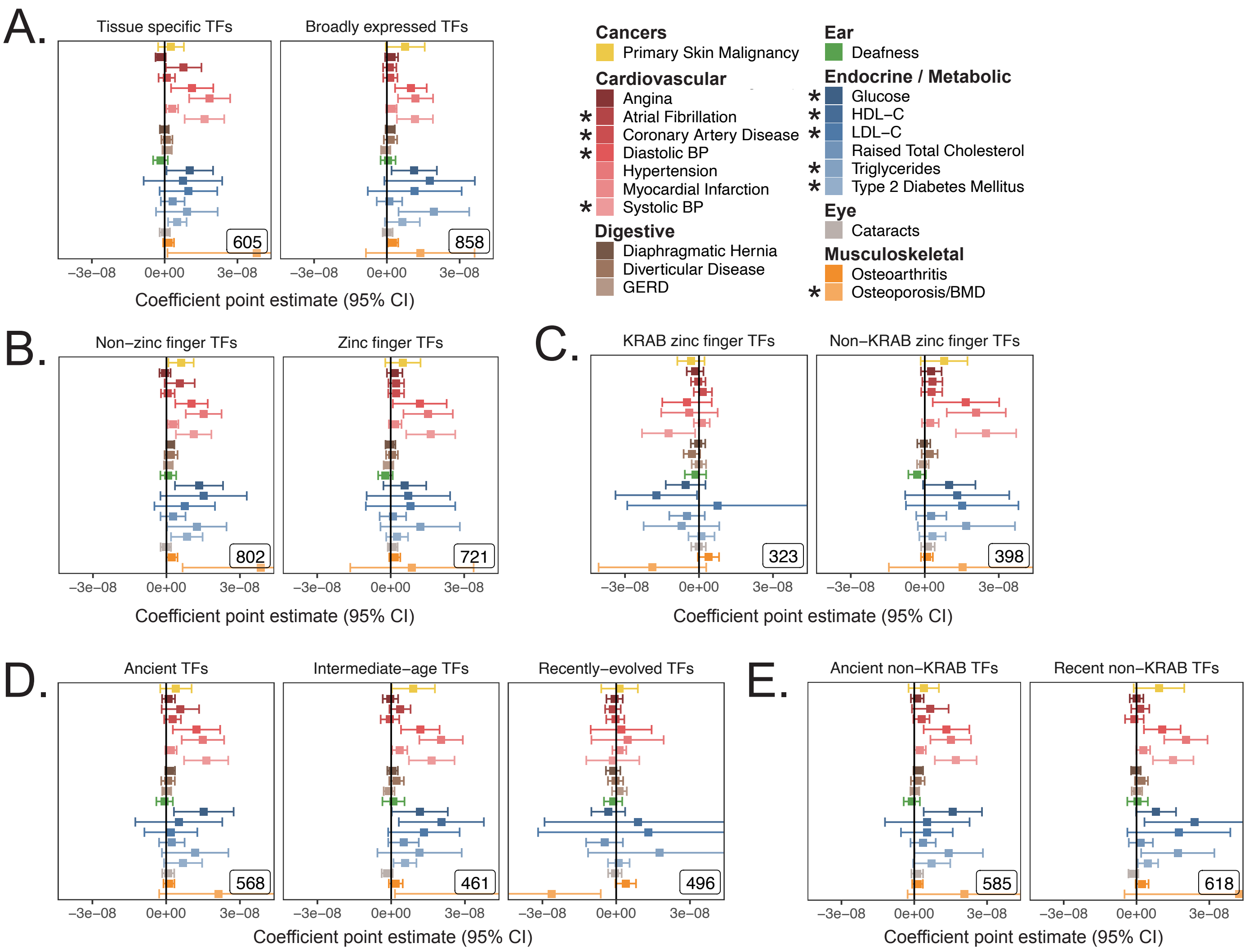


**Figure 3-S7.** S-LDSC enrichment coefficient point estimates in functional subsets of transcription factors corresponding to p-values in **Figure 3-S6**. **A.** Point estimates for heritability enrichment when subdividing TFs based on breadth of tissue expression in GTEx v7 (**Figure 3-S5A**, **3-S5B**). **B.** Point estimates for heritability enrichment when dividing TFs based on the presence of a zinc finger domain. **C.** Point estimates for heritability enrichment when subdividing TFs based on the presence of KRAB domain. **D.** Point estimates for heritability enrichment when dividing TFs into terciles based on evolutionary gene age (see **Figure 3-S5F**). **E.** Point estimates for heritability enrichment when subdividing TFs into old and new age bins after removing KRAB domain zinc finger TFs. Error bars represent 95% CI.

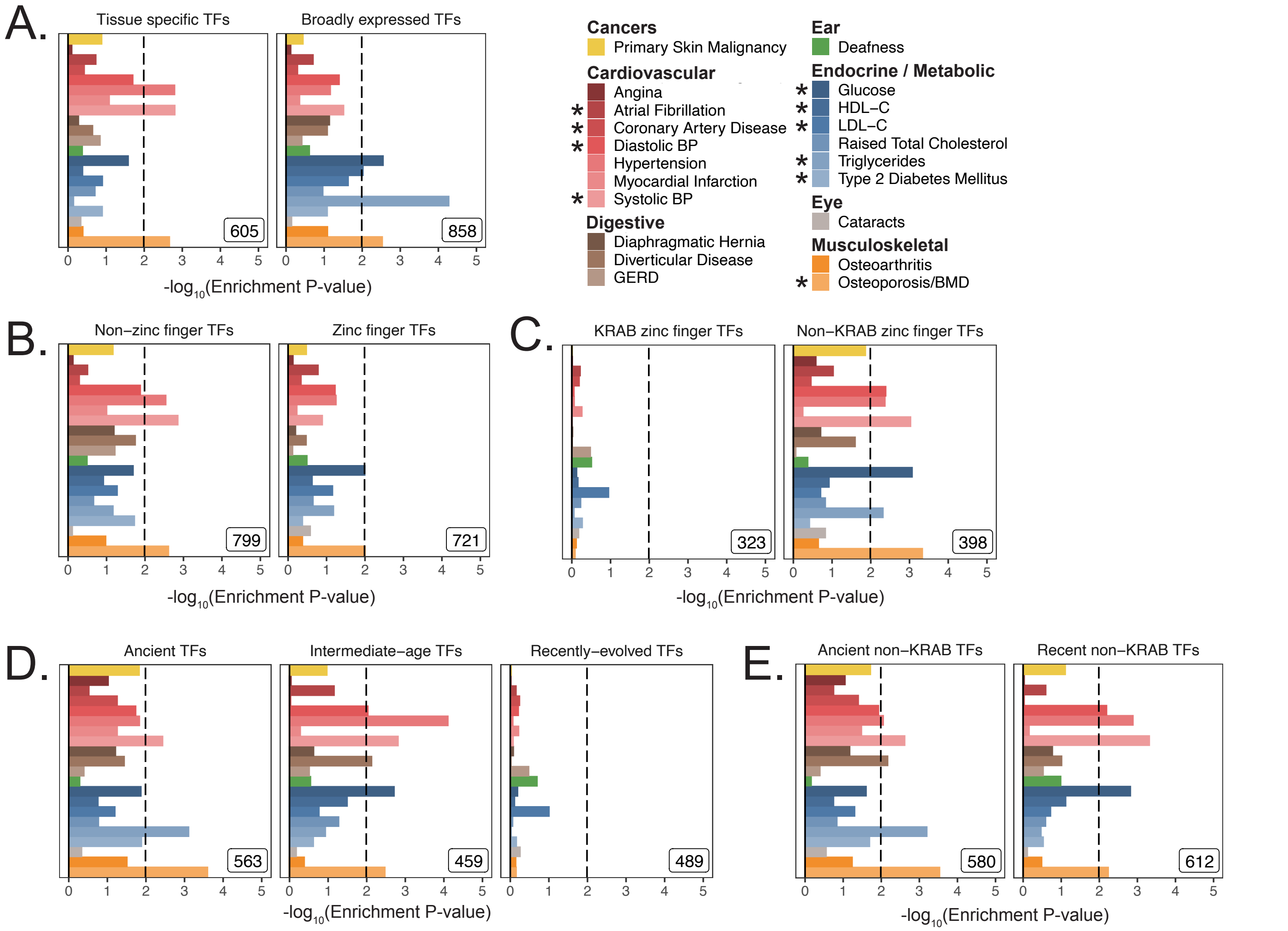


**Figure 3-S8.** Replication of enrichment analysis in functional subsets of transcription factors using MAGMA with a 100kb symmetric gene window. **A.** Enrichment analysis of TFs, subdivided by breadth of tissue expression in GTEx v7 (**Figure 3-S5A**, **3-S5B**). **B.** Enrichment analysis of TFs subdivided by the presence of a zinc finger domain. **C.** Enrichment analysis of zinc finger TFs, subdivided by presence of KRAB domain. **D.** Enrichment results dividing TFs into terciles based on evolutionary gene age (see **Figure 3-S5F**). **E.** Enrichment analysis subdividing TFs into old and new after removing KRAB domain zinc finger TFs. Inset numbers represent gene-set size, black lines represent cutoff at BH FDR < 10%.

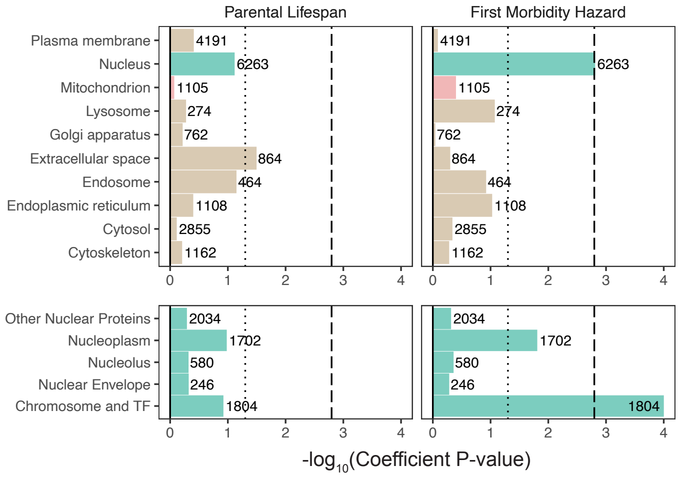


**Figure 4-S1.** Enrichment of organellar proteomes within parental lifespan and healthspan as proxies for aging using S-LDSC. Upper panels represent organelle proteomes; lower panels represent spatial subsets of the nuclear proteome. Numbers atop each bar represent gene-set sizes. Dashed lines represent cutoff at BH FDR < 10%, dotted lines represent nominal $p = 0.05$.

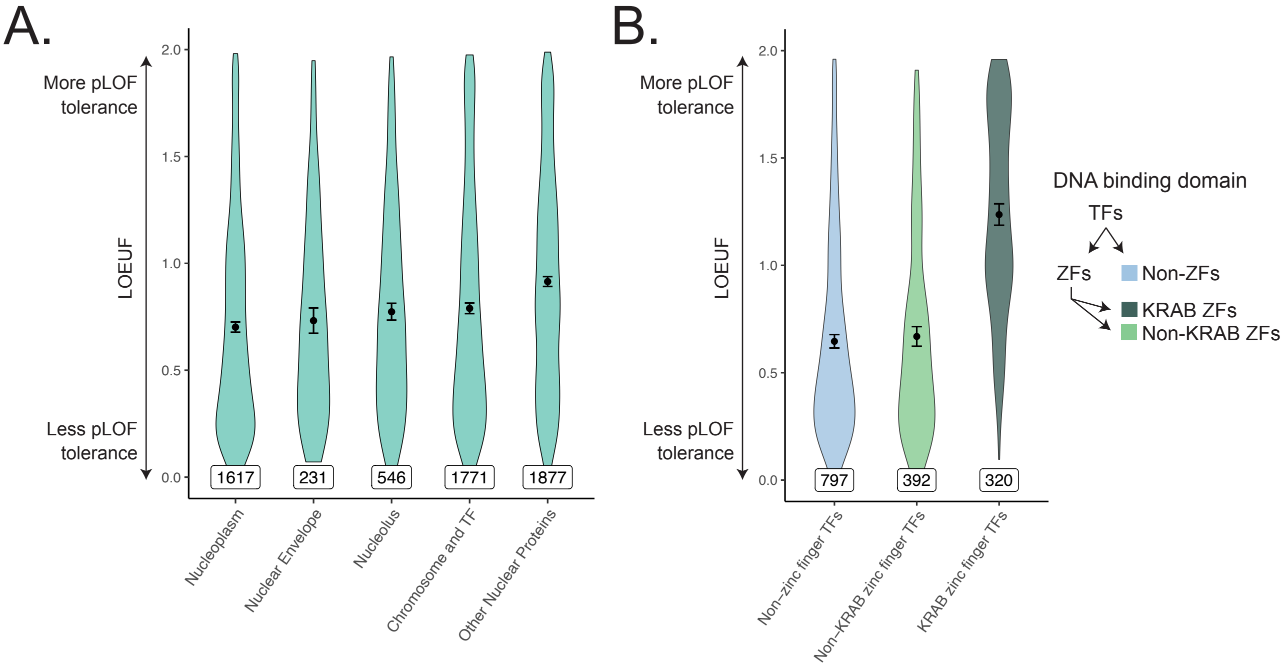


**Figure 5-S1.** Constraint distributions within subdivisions of the nuclear proteome. **A.** Constraint distributions for spatially defined subsets of the nuclear proteome. Gene-sets used are identical to those tested in **Figure 3C**. **B.** Constraint distributions for transcription factor DNA binding domains. Gene-sets used are identical to those tested in **Figure 3-S5**. Black points represent the mean with 95% CI. Inset numbers represent gene-set size.

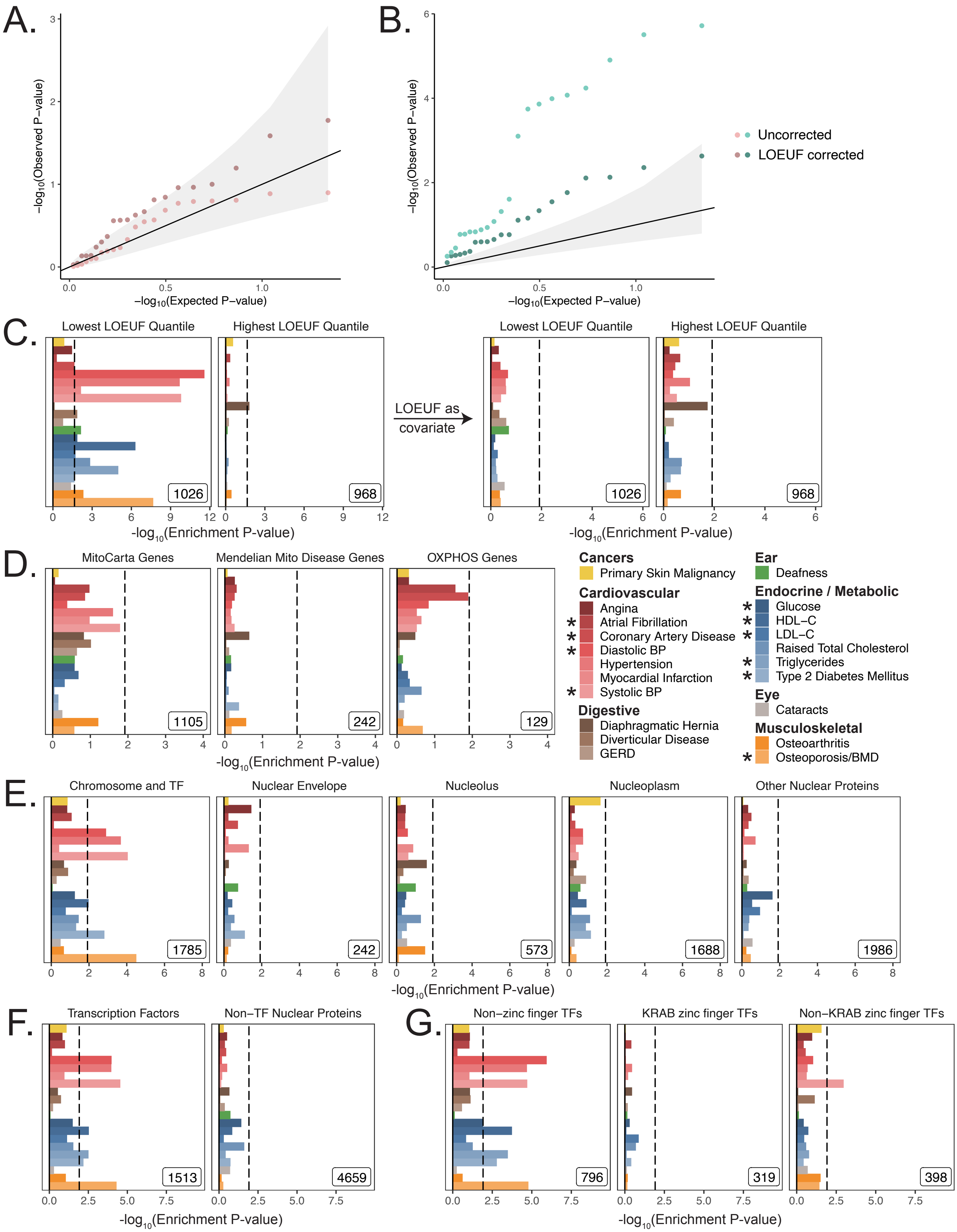


**Figure 5-S2.** Enrichment results across age-related disease in UKB after correcting for constraint using MAGMA with a 5kb up, 1.5kb down window. Quantile-quantile plots showing enrichment across age-associated disease in UKB for **A.** mitochondria-localizing and **B.** nucleus-localizing genes before and after correction for constraint. The central black line represents null p-values following the uniform distribution and the shaded ribbon represents 95% CI. **C.** Enrichment p-values for genes with the lowest and highest LOEUF before and after inclusion of constraint as a covariate in the MAGMA model. **D.** Enrichment p-values for mitochondria-localizing genes. **E.** Enrichment p-values for spatial subsets of the nuclear proteome and **F.** for subsets of the Chromosome and TF category. **G.** Enrichment p-values for TFs subdivided by DNA binding domain. Inset numbers represent gene-set size, black lines are cutoff at BH FDR < 10%.

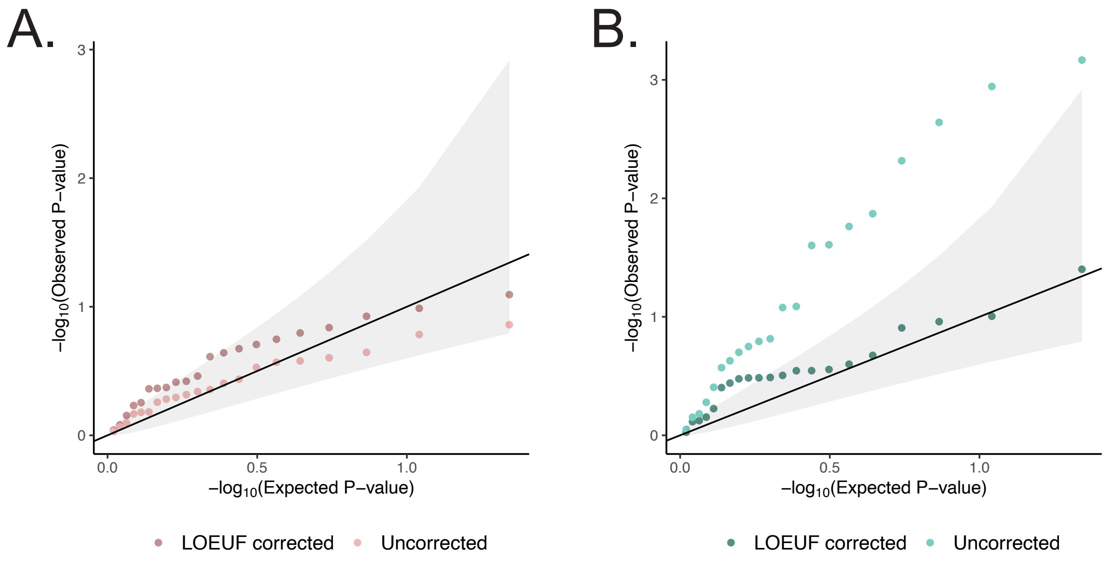


**Figure 5-S3.** Enrichment results across age-related disease in UKB after correcting for gene constraint using MAGMA with a 100kb symmetric gene window. **A.** Quantile-quantile plot highlighting enrichment results for mitochondria-localizing genes across age-associated disease in UKB before and after correcting for constraint. **B.** Quantile-quantile plot highlighting enrichment results for genes comprising the nuclear proteome across age-associated disease in UKB before and after correcting for constraint. The central black line represents expected null p-values following the uniform distribution and the shaded ribbon represents 95% CI.

**Table S1.** Tested traits and sample sizes in UK Biobank and external meta-analyses.

| | | UK Biobank | | | | | | Meta analysis | | | | | |
|---|---|---|---|---|---|---|---|---|---|---|---|---|---|
| Phenotype | Median onset age | Code | N | Cases | Controls | $h^2$ | $h^2_{se}$ | Study | N | Cases | Controls | $h^2$ | $h^2_{se}$ |
| GERD | 50 | K21 | 361194 | 10743 | 350451 | 0.08 | 0.01 | | | | | | |
| Deafness | 52 | 3393 | 219358 | 10942 | 208416 | 0.10 | 0.02 | | | | | | |
| Raised Total Cholesterol | 56 | 20002_1473 | 361141 | 43957 | 317184 | 0.12 | 0.01 | | | | | | |
| HDL-C | 57† | 30760_irnt | 315133 | – | – | 0.22 | 0.01 | Teslovich et al., 2010 | 99900* | – | – | 0.12 | 0.01 |
| LDL-C | 57† | 30780_irnt | 343621 | – | – | 0.10 | 0.01 | Teslovich et al., 2010 | 95454* | – | – | 0.10 | 0.01 |
| Triglycerides | 57† | 30870_irnt | 343992 | – | – | 0.17 | 0.01 | Teslovich et al., 2010 | 96598* | – | – | 0.12 | 0.02 |
| Diastolic BP | 59† | 4079_irnt | 340162 | – | – | 0.14 | 0.01 | ICBP 2011 | 69395* | – | – | 0.14 | 0.02 |
| Hypertension | 59 | 20002_1065 | 361141 | 93560 | 267581 | 0.22 | 0.01 | | | | | | |
| Systolic BP | 59† | 4080_irnt | 340159 | – | – | 0.15 | 0.01 | ICBP 2011 | 69395* | – | – | 0.14 | 0.02 |
| Diaphragmatic Hernia | 60 | K44 | 361194 | 8042 | 353152 | 0.08 | 0.02 | | | | | | |
| Glucose | 61† | 30740_irnt | 314916 | – | – | 0.08 | 0.01 | Manning et al. 2012 | 58074 | – | – | 0.09 | 0.02 |
| Osteoarthritis | 61 | 20002_1465 | 361141 | 30046 | 331095 | 0.07 | 0.01 | | | | | | |
| Type 2 Diabetes Mellitus | 61 | 2443 | 360192 | 17275 | 342917 | 0.21 | 0.02 | Morris et al. 2012 | 69033* | 12171* | 56862* | 0.18 | 0.03 |
| Angina | 64 | 20002_1074 | 361141 | 11370 | 349771 | 0.17 | 0.02 | | | | | | |
| Myocardial Infarction | 65 | I9_CHD | 361194 | 10157 | 351037 | 0.14 | 0.02 | | | | | | |
| Diverticular Disease | 67 | K57 | 361194 | 12662 | 348532 | 0.13 | 0.01 | | | | | | |
| Primary Skin Malignancy | 68 | C3_SKIN | 361194 | 16531 | 344663 | 0.16 | 0.03 | | | | | | |
| Coronary Artery Disease | 70 | I9_IHD | 361194 | 20857 | 340337 | 0.15 | 0.01 | Schunkert et al., 2011 | 86995* | 22233* | 64762* | 0.07 | 0.01 |
| Osteoporosis/BMD | 71† | 3148_irnt | 206496 | – | – | 0.28 | 0.02 | Estrada et al 2012 | 32961 | – | – | 0.26 | 0.03 |
| Atrial Fibrillation | 74 | I48 | 361194 | 6356 | 354838 | 0.14 | 0.03 | Christophersen et al. 2017 | 133073 | 17931 | 115142 | 0.06 | 0.01 |
| Cataracts | 74 | H26 | 361194 | 11306 | 349888 | 0.09 | 0.01 | | | | | | |
| eGFR | 74† | | | | | | | Pattaro et al., 2016 | 133413* | – | – | 0.12 | 0.01 |
| Parkinson's Disease§ | 74 | | | | | | | Nalls et al., 2019 | 482730** | 33674** | 449056** | 0.03 | 0.00 |
| Alzheimer's Disease | 83 | | | | | | | Lambert et al., 2013 | 54162 | 17008 | 37154 | 0.05 | 0.01 |

We only show sample sizes for traits with heritability Z score ≥ 4 as obtained by S-LDSC.
* variant-specific sample sizes used in analysis; displayed values are maximal values reported in associated publication
** variant-specific sample size used in analysis and to obtain displayed maximal values
§ meta-analysis contains samples also part of UKB
† median age of onset for continuous traits obtained from discretizing distribution by Kuan et al. or from surrogate diseased states (T2DM for Glucose; CKD for eGFR)

**Table S2.** Genetic and phenotypic correlation point estimates and standard errors.

**Table S3.** QC calls assigned to mtDNA variants through manual review of cluster plots.

**Table S4.** mtDNA GWAS summary statistics.

**Table S5.** Transcription factor assignment to broadly-expressed vs tissue-specific categories using two sub-region groupings.

**Table S6.** Coefficient point estimates and p-values for all tested gene-sets and all tested traits.

**Table S7.** Coefficient point estimates and p-values for all tested gene-sets for aging phenotypes.

**Supplementary Note**

*Choice of traits with meta-analyses with cohorts separate from UKB*

For ten traits with well-powered UKB GWAS and meta-analyses, we ensured that the meta-analyses used did not incorporate data from UKB thus allowing their use as replication cohorts. Parkinson's disease[1] and Alzheimer's disease[2] were analyzed as part of larger meta-analyses only due to power limitations in UKB and eGFR was assessed only in the tested meta-analysis. In the case of Parkinson's disease a well-powered GWAS was recently performed and included UKB individuals[1]. Given that this trait was not sufficiently powered for analysis in UKB alone, we chose to proceed with summary statistics from this study. Because mtDNA-GWAS could only be performed in UKB (where we had access to individual-level data), we were unable to explicitly test for mtDNA associations with Parkinson's disease, Alzheimer's disease, and eGFR.

*Heritability Z-score threshold selection*

Total SNP heritability Z-score encapsulates variables such as polygenicity, sample size, and underlying disease heritability, all of which influence S-LDSC power[3]. Previous work has indicated that genetic correlation estimates from LD score regression are noisy for total SNP heritability Z-score < 4[4], and total SNP heritability Z-score > 7 has been used as a condition for trait inclusion for S-LDSC[3]. We decided to use a more relaxed cutoff of total SNP heritability Z-score > 4 for two major reasons: First, we used a distinct enrichment methodology, MAGMA, to validate enrichment signatures. To our knowledge, MAGMA does not produce unstable enrichment estimates for traits with moderate heritability Z-score. Second, we also used GWAS data from non-overlapping cohorts, when available, as independent validation for traits tested in UKB. The lower cutoff was sufficient to produce results that largely replicated across methodology and cohort, while allowing for the inclusion of several traits of interest. Further, several traits with heritability Z-score between 4 and 7 show positive control tissue enrichments and substantial enrichment detection power (for example, LDL levels).

*Choice of traits to test in the GWAS Catalog*

We searched the GWAS Catalog phenotypes to identify age-related traits. We manually identified 30 phenotypes that matched our 24 age-related traits (**Figure 2-S1A**). This list differs from our full list of age-related traits for two reasons: (1) not all 24 age-related traits had a sufficient number of associated genes for analysis, and (2) in several cases multiple phenotypes listed in the GWAS catalog matched our age-related traits (e.g., 'Cholesterol, total' and 'Total cholesterol levels'); we tested these separately.

*Investigation of mitochondria-relevant transcription factors in the GWAS Catalog*

We tested if any of eight TFs known to regulate mitochondrial function – *TFAM, GABPA, GABPB1, ESRRA, YY1, NRF1, PPARGC1A, and PPARGC1B* – were the nearest gene to any genome-wide significant variants listed for age-related traits in the GWAS Catalog. We tested the same traits we used for enrichment analysis of the MitoCarta genes in the GWAS Catalog (**Figure 2-S1**) and did not find any signal for 29/30 tested phenotypes. We did find that TFAM was one of the nearest genes for heel bone mineral density, however we note that there are a total of 1496 unique mapped nearest genes for this trait. Further, we tested mitochondria-localizing genes for enrichment in GWAS for heel bone mineral density (3148_irnt) in UKB and found no evidence of enrichment (**Figure 2**).

*Choice of enrichment method*

In this study, we leveraged several enrichment methods to ensure robustness to methodology. We used Fisher's exact test in a first-pass analysis of enrichment of GWAS signal in the GWAS Catalog. While this provides a useful preview of the enrichment landscape across published GWAS, this suffers from numerous limitations, including the usage of only genome-wide significant SNPs, the treatment of each variant as equally likely to contribute to GWAS signal under the null, and an inability to easily control for covariates such a gene length, among others. As such, we used two different methods, MAGMA and S-LDSC, to test for GWAS enrichment among our gene-sets while resolving these confounders and reducing the likelihood of model misspecification. We used S-LDSC to test for heritability enrichment within specified variants controlling for 53 functional categories including DNase hypersensitivity sites, H3K4Me sites, and coding regions. MAGMA uses a variation of Fisher's method to obtain gene-level test statistics and test for gene-set enrichment controlling for LD structure, and when performing the gene-set enrichment testing we controlled for gene length, inverse MAC, and SNP density. We also used tissue-specific enrichments as positive controls to ensure that the methods we used were working properly.

Notably, when running MAGMA on our age-associated traits on meta-analyses with a 100kb window, we were unable to find tissue-specific enrichments (**Figure 2-S5B**). Given that S-LDSC and MAGMA with a 5kb up and 1.5kb down was able to find these enrichments in the selected meta-analyses (**Figure 2-S2B, 2-S5A**) and that we observe reduced power for enrichments among meta-analyses relative to UKB (**Figure 2-S6H**), we attributed the lack of tissue enrichments using MAGMA at 100kb in meta-analyses to a lack of power. Indeed, MAGMA with a 100kb symmetric window was able to find enrichments in UKB **(Figure 2-S4B)**. Thus, we did not test any other gene-sets among meta-analyses using MAGMA with a 100kb window.

*Choice of control genes for gene-based tests*

For all gene-based analyses we aimed to perform a competitive analysis, testing if our genes of interest explained more trait heritability than comparable loci elsewhere in the genome. For our positive control tests and power analyses leveraging the set of highest expressed tissue-specific genes, we controlled for the set of genes across which t-statistics were computed (~25,000 genes); namely all genes that had at least four samples in GTEx with 1 or more counts-per-million[5]. All of our non-tissue gene-sets (e.g., MitoCarta genes, organelle-localizing genes) were subsets of the set of protein-coding genes, so we controlled for the set of protein-coding genes for these analyses (~19,000 genes). For S-LDSC, this involved including the respective control gene-set annotation atop the baseline model; for MAGMA, this involved defining gene location file based on the control gene-set such that the space of genes considered was restricted to the genes to be controlled for.

*Power analysis of gene-based tests*

To verify the power of S-LDSC and MAGMA in our selected traits, we sub-sampled each of ten positive control tissue-trait pairs. We subsampled the set of tissue-expressed genes for each of the six selected tissues at various gene-set sizes and empirically assessed the number of trials in which significant enrichment was detected, giving us an estimate of power, or $\Pr(reject \mid alternative)$. All tissue enrichments were originally performed with 2485 genes (**Methods**); as such we conducted subsampling trials with 1523, 1105, 800, and 350 genes to assess power throughout our study. Because LD score computations are very computationally intensive, we generated 50 random subsamples per gene-set size-tissue pair ensuring that each sample contained a proportional number of genes per chromosome to the original tissue expressed gene-set. We mapped variants to genes and computed LD scores per-chromosome for each annotation (**Methods**). For each gene-set size and tissue (24 gene-set size-tissue pairs), we generated 1000 sets of LD scores by shuffling LD scores computed per chromosome, effectively generating 1000 random tissue gene-set subsamples for each gene-set size-tissue pair. We subsequently used S-LDSC to test for enrichment for each of the 1000 tissue gene-set subsamples in the aforementioned selected traits for each gene-set size, resulting in 240,000 regressions atop the baseline model as performed for the tissue enrichments in the usual way (**Methods**). The gene-sets generated for use with S-LDSC (1000 per gene-set size-tissue pair) were also exported for analysis using MAGMA with the same competitive analysis performed for the tissue-enrichment analysis (**Methods**).

To characterize the power differential between UKB and meta-analyses, we tested the subset of the tissue-trait pairs tested in UKB that showed enrichment in the corresponding meta-analysis with either S-LDSC (**Figure 2-S2B**) or MAGMA (**Figure 2-S4B**). This resulted in an assessment of power among meta-analyses for liver-TG, liver-LDL, liver-HDL, visceral adipose-HDL, atrial appendage-atrial fibrillation, pancreas-T2D, and pancreas-glucose. We tested the same gene-sets tested with S-LDSC and MAGMA (1000 per gene-set size-tissue pair) in UKB using both S-LDSC and MAGMA in the usual way (**Methods**).

As expected, we noted that power was a function of both enrichment effect size and gene-set size for S-LDSC and MAGMA (**Figure 2-S6A–2-S6F**). While we observed lower power across most tested traits among meta-analyses when compared to UKB, power was acceptable among the meta-analyses for high effect size enrichments for gene-sets with 1105 genes (**Figure 2-S6G, 2-S6H**).

*Choice of gene-sets to test for replication among meta-analyses*
Because our power analyses showed a substantial reduction of power for tested meta-analyses relative to UKB (**Figure 2-S6I, 2-S6J**), we tested only a subset of all tested gene-sets for replication among meta-analyses. Namely, we sought to test replication of the two major organelle-based results in this study: (1) the lack of enrichment of mitochondria-localizing genes across age-related disease, and (2) the enrichment of chromosome and TF genes, with subsequent enrichment of the TFs alone.

*Choice of tissues to include for multi-tissue analyses*
An assumption key to several statistics used for the eQTL and TF breadth of expression analyses ($\tau_x$, $N_x^{eQTL}$, $N_x^{express}$) is that different tissues are not overrepresented in the set of tissues assessed. This assumption breaks down in GTEx, where the brain, artery, and esophagus were sampled in multiple sub-regions and the skin, cervix, colon, and adipose tissue were sampled in two sub-regions. We selected specific sub-regions as shown in **Figure 3-S5B** manually as expression profiles within sub-regions tend to be far more similar than profiles between sub-regions. To test robustness, we selected an alternate set of tissues within each class (brain frontal cortex (ba9), artery tibial, esophagus muscularis, skin sun exposed (lower leg), colon transverse, and adipose subcutaneous) and repeated our analyses. For our eQTL analysis, we find results that are very similar using this alternate set of tissues as expected **(Figure 2-S8C**). Further, we found that with the cutoff of $\tau = 0.76$ for a tissue specific gene, only 32 of the 1,463 tested TFs would be classified differently (**Table S5**). Using our original choice of tissues, we find that 605 TFs are tissue specific (Lambert et al. report 542 tissue-specific TFs), that 75% of homeodomain containing TFs are tissue specific (Lambert et al. report 82%), and that 18.6% of KRAB ZF TFs are tissue specific (Lambert et al. report 12%). Thus, our results using our choice of tissues are robust to the specific choice of tissue sub-region within a tissue region and are in good agreement with previously reported tissue-specific expression annotations.

*Model selection for eQTL analyses*
To understand if genetic variation near genes localizing to a given organelle were abnormally unlikely to produce downstream biological consequences, we turned to cis-eQTLs. Because most genes have a measured cis-eQTL in at least one tissue (**Figure 2-S8A**), we constructed a model to test if genes localizing to a given organelle had significant cis-eQTLs in more or less tissues than other protein-coding genes. We included several covariates to minimize the risk of confounding from first principles (**Methods**). We corrected for $gene\ length$ and $\log_{10}(gene\ length)$ as we expected that higher number of SNPs in longer genes would increase the probability of eQTL detection; $N_x^{express}$ as we suspected that genes would have detectable eQTLs at most in tissues where they were expressed; and $\tau_x$ as we expected that broadly

expressed genes would be more likely to have cis-eQTLs detected in more tissues. Upon model fitting, we observed that all coefficients were significantly different from 0.

*Manual variant QC for mtDNA-GWAS*

We used two strategies to manually review the variants that made it through automated variant QC filters (**Methods**). First, we visually reviewed fluorescence cluster plots for each mtDNA variant to ensure that our variant calls were accurate (**Methods**). We visually categorized each variant into 5 categories: clear pass, batch concern, off target variant (OTV) concern, resolution concern, and misclustering (**Table S3**), removing 19 variants from further analysis due to cluster plot abnormalities. Second, we computed the mtDNA LD matrix finding no evidence of distance-dependent LD on the mtDNA (**Figure 2-S9A**) as observed previously[6].

*Minor allele frequency filters for mtDNA-GWAS*

We used two variant frequency filters to ensure that our regression test statistics were well-behaved (**Methods**). For continuous traits, we included only variants that had at least 20 individuals with an alternate genotype. For binary traits, we implemented a per-trait and per-variant filter by computing the proportion of individuals with an alternate genotype required such that, under null expectation, there would be 20 cases with an alternate genotype. This filter has been shown to eliminate false positive associations by eliminating low MAC variants for rare traits, in which highly imprecise allele frequency estimates can exert high leverage on test statistics[7]. This was operationalized as a MAF cutoff as there are by definition no heterozygotes on the mitochondrial DNA, such that for each trait we included only variants that satisfied $MAF \geq 20/\min(CaseSampleSize, ControlSampleSize)$. The sample size estimates were dependent on the variant being assessed as certain variants had distinct missingness patterns due to measurement on a particular genotype array used for only a subset of the cohort (**Methods**). In total, we tested up to 213 variants per phenotype, assessing a total of 4337 variant-phenotype pairs.

*Enrichment analysis of Parkinson's Disease*

Of course, much interest lies around characterizing the involvement of mitochondrial dysfunction in PD[8–11]. We find no evidence of heritability enrichment among MitoCarta genes in a recent PD GWAS[1] (**Figure 2D**). Due to power limitations, we were unable to assess mtDNA associations with PD (**Supplementary note**), though to our knowledge, broadly reproducible associations between inherited mtDNA variants and PD have yet to be reported[12,13].

*Interpretation of heritability explained by organellar gene-sets*

For the sets of genes corresponding to organellar proteomes, we highlight the substantial amount of SNP-heritability explained by variants in or near genes contributing to the nuclear proteome. It is notable that all organelles show $prop\ h^2_{SNP}\ / prop\ SNP > 1$ (**Figure 3-S1**). We believe that this is because of other properties of the SNPs near organelle-localizing genes, namely that all selected SNPs are near protein coding genes. SNPs in protein coding regions are known to be enriched for heritability[3], and indeed when we explicitly model these potentially confounding functional SNP annotations (DNase hypersensitivity sites, H3K4Me sites, coding regions; **Methods**) only the enrichment among variants near nucleus-localizing genes persists.

*Overlap analysis of subsets of the nuclear proteome*

We performed pairwise overlap analysis for our five final sub-nuclear compartments (Nucleoplasm, Chromosome and TF, Nucleolus, Nuclear Envelope, Other Nuclear Proteins), finding that virtually all pairs showed an overlap of less than 5% (with an exception for the nucleolus, ~13% of which was also represented in chromosome and TF). S-LDSC and MAGMA were used to test for enrichment across the UKB age-related traits for these gene-sets as performed previously for the organelle analysis.

*GWAS enrichments of functional subdivisions of the class of TFs*

We further subdivided the TFs based on breadth of expression in human tissues, DNA binding domain (DBD), and gene age (**Methods**). We found a similar pattern of enrichment for tissue-specific TFs and broadly expressed TFs (**Figure 3-S5C, 3-S6A, 3-S7A**). However, upon stratification by the three largest categories of TF DBD[14], we found that non-zinc finger TFs showed enrichment for many age-related traits (**Figure 3-S5D, 3-S6B, 3-S7B, 3-S8B**), while the KRAB domain-containing zinc fingers (KRAB ZFs), were largely devoid of enrichment even compared to non-KRAB ZFs (**Figure 3-S5E, 3-S6C, 3-S7C, 3-S8C**). While our power analysis suggests sufficient power only for high effect sizes at ~350 genes, we note that (1) the KRAB ZFs and non-KRAB ZFs have similar gene-set sizes and (2) S-LDSC coefficient point estimates are systematically much higher for non-KRAB ZFs than for KRAB ZFs (**Figure 3-S7C**). Notably, while we initially observed enrichment only for ancient and intermediate-age TFs but not recently-evolved TFs (**Figure 3-S5G, 3-S6D, 3-S7D, 3-S8D**), we find that old and recent non-KRAB TFs (**Figure 3-S5C**) showed similar enrichment profiles (**Figure 3-S5I, 3-S6E, 3-S7E, 3-S8E**), suggesting that the lack of signal among recent TFs was likely attributable to the KRAB domain containing ZFs which are predominantly recently-evolved (**Figure 3-S5H**).

*Age-related disease GWAS enrichment with constraint as a covariate*

We wanted to assess if our observed enrichment results persist after explicitly accounting for any variance explained by the degree of constraint. We used MAGMA and included LOEUF as a covariate in the gene-set enrichment analysis model (**Methods**), finding that the LOEUF correction did not substantially impact MitoCarta gene enrichment (**Figure 5-S2A, 5-S3A**) but did reduce the degree of enrichment seen for nucleus-localizing genes (**Figure 5-S2B, 5-S3B**). We continue observing enrichment for the TFs across several age-related diseases (**Figure 5-S2E, 5-S2F**) with a similar pattern of enrichment in non-ZF TFs and non-KRAB ZFs (**Figure 5-S2G**) to that seen with the original model (**Figure 3-S5D, 3-S5E**). Thus, while constraint explains a substantial component of the enrichment observed for the TFs among age-related diseases, an enrichment signal persists after accounting for LOEUF.

1. Nalls MA, Blauwendraat C, Vallerga CL, et al. Identification of novel risk loci, causal insights, and heritable risk for Parkinson's disease: a meta-analysis of genome-wide association studies. *Lancet Neurol*. 2019;18(12):1091-1102. doi:10.1016/S1474-4422(19)30320-5
2. Lambert JC, Ibrahim-Verbaas CA, Harold D, et al. Meta-analysis of 74,046 individuals identifies 11 new susceptibility loci for Alzheimer's disease. *Nat Genet*. 2013;45(12):1452-1458. doi:10.1038/ng.2802
3. Finucane HK, Bulik-Sullivan B, Gusev A, et al. Partitioning heritability by functional annotation using genome-wide association summary statistics. *Nat Genet*. 2015;47(11):1228-1235. doi:10.1038/ng.3404
4. Bulik-Sullivan B, Finucane HK, Anttila V, et al. An atlas of genetic correlations across human diseases and traits. *Nat Genet*. 2015;47(11):1236-1241. doi:10.1038/ng.3406
5. Finucane HK, Reshef YA, Anttila V, et al. Heritability enrichment of specifically expressed genes identifies disease-relevant tissues and cell types. *Nat Genet*. 2018;50(4):621-629. doi:10.1038/s41588-018-0081-4
6. Yamamoto K, Sakaue S, Matsuda K, et al. Genetic and phenotypic landscape of the mitochondrial genome in the Japanese population. *Commun Biol*. 2020;3(1):104. doi:10.1038/s42003-020-0812-9
7. Howrigan D, Abbot L, Churchhouse C, Palmer DS. Details and considerations of the UK Biobank GWAS. Neale lab blog. http://www.nealelab.is/blog/2017/9/11/details-and-considerations-of-the-uk-biobank-gwas. Published 2017.
8. Nguyen M, Wong YC, Ysselstein D, Severino A, Krainc D. Synaptic, Mitochondrial, and Lysosomal Dysfunction in Parkinson's Disease. *Trends Neurosci*. 2019;42(2):140-149. doi:10.1016/j.tins.2018.11.001
9. Grünewald A, Kumar KR, Sue CM. New insights into the complex role of mitochondria in Parkinson's disease. *Prog Neurobiol*. 2019;177(April 2018):73-93. doi:10.1016/j.pneurobio.2018.09.003
10. Abou-Sleiman PM, Muqit MMK, Wood NW. Expanding insights of mitochondrial dysfunction in Parkinson's disease. *Nat Rev Neurosci*. 2006;7(3):207-219. doi:10.1038/nrn1868
11. Ge P, Dawson VL, Dawson TM. PINK1 and Parkin mitochondrial quality control: A source of regional vulnerability in Parkinson's disease. *Mol Neurodegener*. 2020;15(1):1-18. doi:10.1186/s13024-020-00367-7
12. Bose A, Beal MF. Mitochondrial dysfunction in Parkinson's disease. *J Neurochem*. 2016;139:216-231. doi:10.1111/jnc.13731
13. Müller-Nedebock AC, Brennan RR, Venter M, et al. The unresolved role of mitochondrial DNA in Parkinson's disease: An overview of published studies, their limitations, and future prospects. *Neurochem Int*. 2019;129(April):104495. doi:10.1016/j.neuint.2019.104495
14. Lambert SA, Jolma A, Campitelli LF, et al. The Human Transcription Factors. *Cell*. 2018;172(4):650-665. doi:10.1016/j.cell.2018.01.029
